# Early lineage divergence segregates sensory and non-sensory thalamic circuits

**DOI:** 10.1101/2025.07.17.665342

**Authors:** Lorenzo Puche-Aroca, Belén Andrés-Bayón, Emily S. Wilson, Awais Javed, Aurelia Torregrosa-Mira, José P. López-Atalaya, Guillermina López-Bendito

**Affiliations:** Institute of Neurosciences, Miguel Hernández University - Consejo Superior de Investigaciones Científicas (UMH-CSIC), San Juan de Alicante, Alicante, Spain; Department of Basic Neuroscience, University of Geneva, Geneva, Switzerland

**Author notes:** Correspondence (J.L-A); (G.L-B).

**Keywords:** Development, sensory circuits, motor circuits, thalamus, neural progenitors, fate specification, DNA barcoding lineage tracing, multiomics, spatial transcriptomics

## Abstract

The thalamus integrates sensory, motor, and associative functions via distinct nuclei, yet the developmental principles driving their specification remains unclear. Here, we generate a spatiotemporal single-cell multiomic and spatial transcriptomic atlas of the embryonic mouse thalamus, combined with barcoding-based lineage tracing. We identify two major glutamatergic lineages—sensory and motor/associative—each governed by different gene regulatory programs and temporal dynamics. These lineages arise from spatially and molecularly isolated progenitors, revealing an early segregation of functional identities. Within the sensory lineage, we resolve two discrete trajectories leading to first-order visual and somatosensory neurons, both emerging from a shared higher-order default state. Using *in silico* predictions and *in vivo* perturbations, we identify *Sp9* as a key regulator of visual thalamic fate. Together, our findings define the molecular architecture and developmental trajectories that underpin thalamic modality specification, providing a framework to understand how functional circuits emerge in the mammalian brain.

## Introduction

The thalamus serves as a central hub for sensory, motor and associative integration, comprising anatomical and functionally distinct nuclei that mediate both peripheral inputs and reciprocal communication with the cortex^1,2^. These nuclei are broadly classified into first-order (FO) sensory nuclei (e.g., dLG, VPM, MGv), which rely peripheral signals to primary cortical areas; higher-order (HO) sensory nuclei (e.g., LP, Po, MGd), which participate in cortico-thalamo-cortical loops that support integrative sensory processing; and motor nuclei (e.g. VA, VL, VM) and associative nuclei (e.g. CM, MD), which contribute to motor planning and executive functions^3–7^. Each nucleus is characterised by distinct anatomical projections, electrophysiological properties, and gene expression profiles^8–10^. However, the developmental mechanisms that give rise to this functional and molecular diversity remain elusive.

Thalamic excitatory neurons originate from progenitor cell located in the caudal thalamic neuroepithelium during embryogenesis^11,12^. While prior studies have defined broad neurogenic timelines and identified some fate restrictions within sensory thalamic populations^13–17^, the molecular and temporal mechanisms that direct the specification of divergent thalamic lineages and their functional modalities are poorly understood. Here, we generated a comprehensive spatiotemporal single-cell multiomic and spatial transcriptomic atlas of the developing mouse thalamus, encompassing key stages of neurogenesis and lineage commitment. We identified two major glutamatergic lineages with different temporal dynamics that give rise to sensory and non-sensory thalamic classes. Within the sensory lineage, we resolved two divergent trajectories leading to FO visual dLG and somatosensory VPM nuclei, both emerging from a shared prenatal HO-like molecular state. Finally, we identified Sp9 as a lineage-specific regulator of visual thalamic identity. These findings establish a developmental framework for thalamic specification and reveal how early lineage gene programs shape the emergence of functional circuits in the mammalian brain.

## Results

### Ontogeny of thalamic neurons revealed by single-cell multiomic profiling

To dissect the transcriptional and epigenetic programs underlying thalamic neurogenesis, we combined *in vivo* pulse-labelling of isochronic cell cohorts (FlashTag, FT)^18^ with single-cell multiomic profiling (scRNA-seq and scATAC-seq) across embryonic development. FT was injected into the third ventricle of mouse embryos at gestational day (E)11.5, E14.5, and E16.5, and labelled cells were isolated by fluorescence-activated cell sorting (FACS) 12 hours later. This approach enabled temporal capture and profiling of mitotic progenitors and their immediate postmitotic progeny as they migrate through the developing thalamus (Fig. 1a–d).

**Fig. 1.**
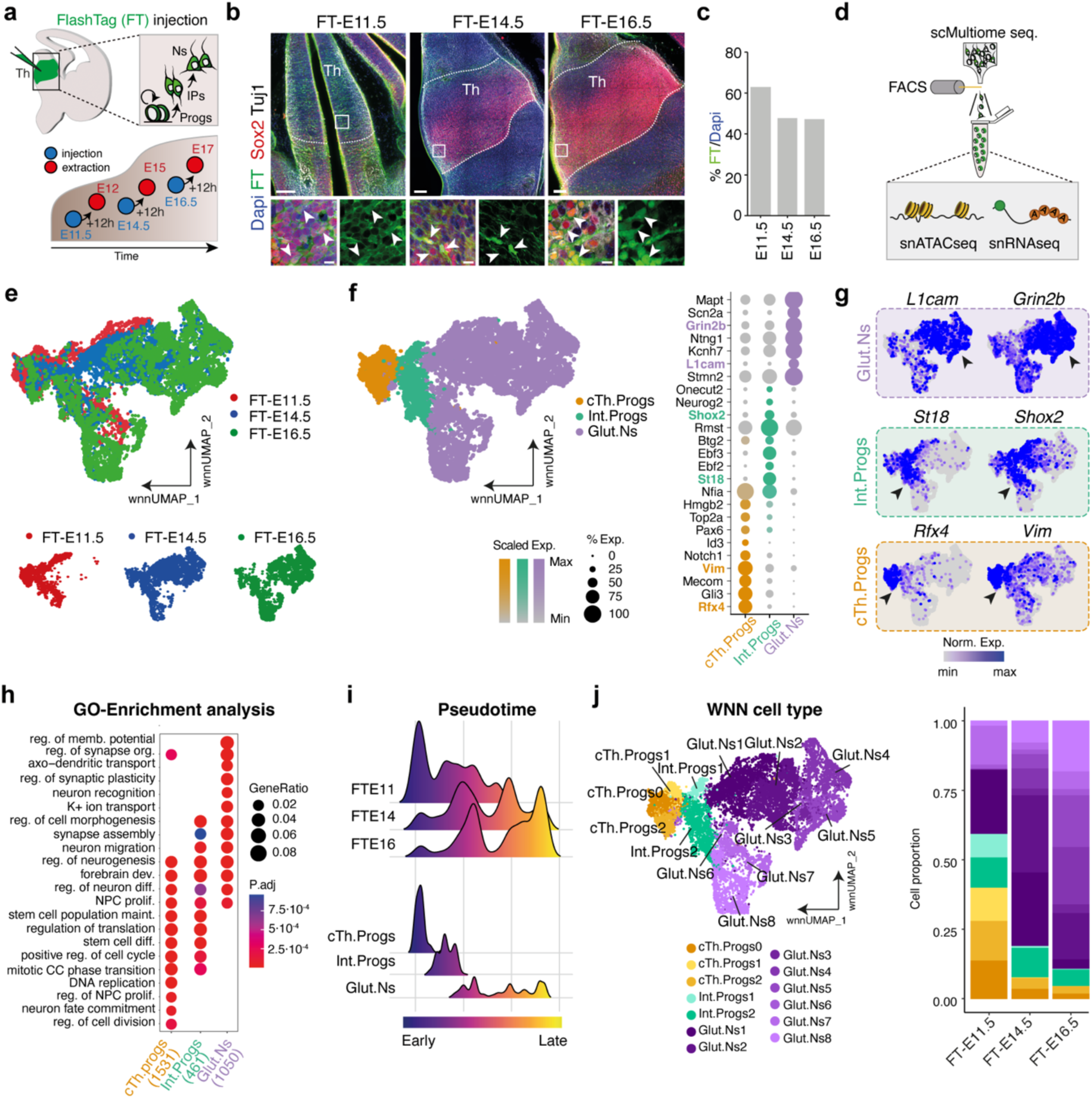
A multimodal atlas of thalamic glutamatergic neuron development. **a**, Schematic of the experimental pipeline combining *in vivo* FlashTag (FT) pulse-labeling with single-cell multiomic profiling. **b**, Coronal sections of the embryonic mouse thalamus showing FT-labeled apical progenitors in M-phase following FT injection at E11.5, E14.5, and E16.5. Labeled progenitors and their daughter cells were isolated 12 h post-injection. Insets: super-resolution images highlighting FT⁺Sox2⁺ cells (arrowheads) at each stage. **c**, Quantification of FT signal intensity relative to DAPI across developmental timepoints in the whole thalamus. **d**, Schematic summarizing the workflow for single-nuclei transcriptome (snRNAseq) and chromatin accessibility profiling (snATACseq) after cell sorting (FACS). **e**, Weighted Nearest Neighbour (WNN) UMAP of paired scRNA-seq and scATAC-seq profiles from glutamatergic thalamic cells at each developmental stage (E11.5+12 h, E14.5+12 h, E16.5+12 h); bottom: the same data arranged by pseudotime. **f**, Left: UMAP identifying three major cell classes. Right: dot plot of representative marker genes defining each class. **g**, Feature plots showing expression of selected markers from (f). **h**, Gene Ontology (GO) enrichment analysis of biological processes by cell type. **i**, Pseudotime trajectory density plots grouped by developmental stage (top) and cell type (bottom). **j**, Left: multimodal UMAP of annotated cell types. Right: proportions of each cell type across E11.5, E14.5, and E16.5. Scale bars, b 100 μm, (insets) 10 μm.

We generated 25,477 high-quality single-cell transcriptomic and epigenomic profiles, capturing all major cell types of the developing thalamus (Extended Data Fig. 1,2). To resolve the earliest stages of lineage specification, we focused on the E11.5 dataset, which reflects the onset of progenitor heterogeneity^19^. Proliferating populations were identified based on cell cycle phase, with actively cycling cells (S and G2/M) forming transcriptionally distinct clusters (Prog1-Prog4) (Extended Data Fig. 3a,b). We identified a pan-radial glia-like progenitor (Prog1; *Id3*^14^), and three distinct progenitor domains: Prog2, enriched in *Tcf7l2*, represents caudal progenitors^14,17,20^; Prog3, marked by *Nkx2.2* and *Arx*, corresponds to rostral progenitors^15,17^, and Prog4 that defined a distinct subset with expression of *Shh, Pitx2, Sim2*, consistent with progenitors of the zona limitans intrathalamica (ZLI)^15,17,21^ (Extended Data Fig. 3c,d).

To assign spatial identity to these molecularly defined progenitor populations, we leveraged a published genome-wide spatial transcriptomic atlas of E9.5 mouse forebrain^22^. This mapping confirmed that Prog1-Prog4 are spatially segregated along the rostro-caudal axis and are located surrounding their derived neural domains, supporting the classical “outside-in” model of thalamic neurogenesis^15,23^ (Extended Data Fig. 3e–g). Furthermore, these progenitors displayed early commitment, generating caudal Prog2 glutamatergic thalamic neurons and rostral Prog3 progenitors fated to produce GABAergic neurons (Extended Data Fig. 3h). Together, these analyses reveal a transcriptionally and spatially organised progenitor landscape that emerges at the onset of thalamic neurogenesis (Extended Data Fig. 3i).

Clustering and trajectory analysis of the caudal progenitor (Prog2) and its progeny across developmental timepoints revealed a continuous differentiation trajectory from neural stem cells to postmitotic neurons, via intermediate progenitor stages (Fig. 1e–g, Extended Data Fig. 4). We identified three major cell classes: caudal thalamic progenitors (cTh.Progs) expressing marked genes such as *Vim* and *Rfx4*^16,24^, intermediate progenitors (Int.Progs) marked by *St18* and *Shox2*^16^, and postmitotic glutamatergic neurons (Glut.Ns) enriched in *L1cam,* and *Grin2b*^16,25,26^ (Fig. 1f,g). Gene ontology (GO) analysis within the joint RNA-ATAC embedding confirmed functional distinctions among these classes, cTh.Progs were enriched for cell cycle and DNA replication genes, Int.Progs exhibited both proliferative and neurogenic signatures, and Glut.Ns displayed upregulated genes associated with synaptic transmission and membrane excitability (Fig. 1h). Pseudotime reconstruction mapped a coherent developmental continuum, extending from early progenitors at E12 to mature glutamatergic neurons at E17 (Fig. 1i). Further clustering analysis at higher resolution revealed three subclasses of cTh.Progs (cTh.Progs0, cTh.Progs1 and cTh.Progs2), two Int.Progs (Int.Progs1 and Int.Progs2), and eight transcriptionally distinct Glut.Ns subtypes, including four terminal populations (Glut.Ns4, Glut.Ns5, Glut.Ns7 and Glut.Ns8) (Fig. 1j). Together, these data define a continuous trajectory from spatially distinct progenitors to discrete glutamatergic neuron subtypes derived from caudal thalamic lineages.

### Time-encoded lineages diverge into sensory and non-sensory thalamic fates

We next asked whether these diverse Glut.Ns subpopulations arise from a common developmental origin or represent divergent lineages. Trajectory reconstruction of our multiomic atlas identified two major glutamatergic neuronal lineages, Lineage 1 and Lineage 2, emerging from caudal thalamic progenitors cTh.Progs1 and cTh.Progs2, respectively (Fig. 2a). Lineage 1 (blue) followed an accelerated differentiation trajectory relative to Lineage 2 (purple), and encompassed Glut.Ns4 and Glut.Ns5 (hereafter Glut.NsLin1). Lineage 2 comprised Glut.Ns7 and Glut.Ns8 (hereafter Glut.NsLin2) (Fig. 2a,b, Extended Data Fig. 5a,b).

**Fig. 2.**
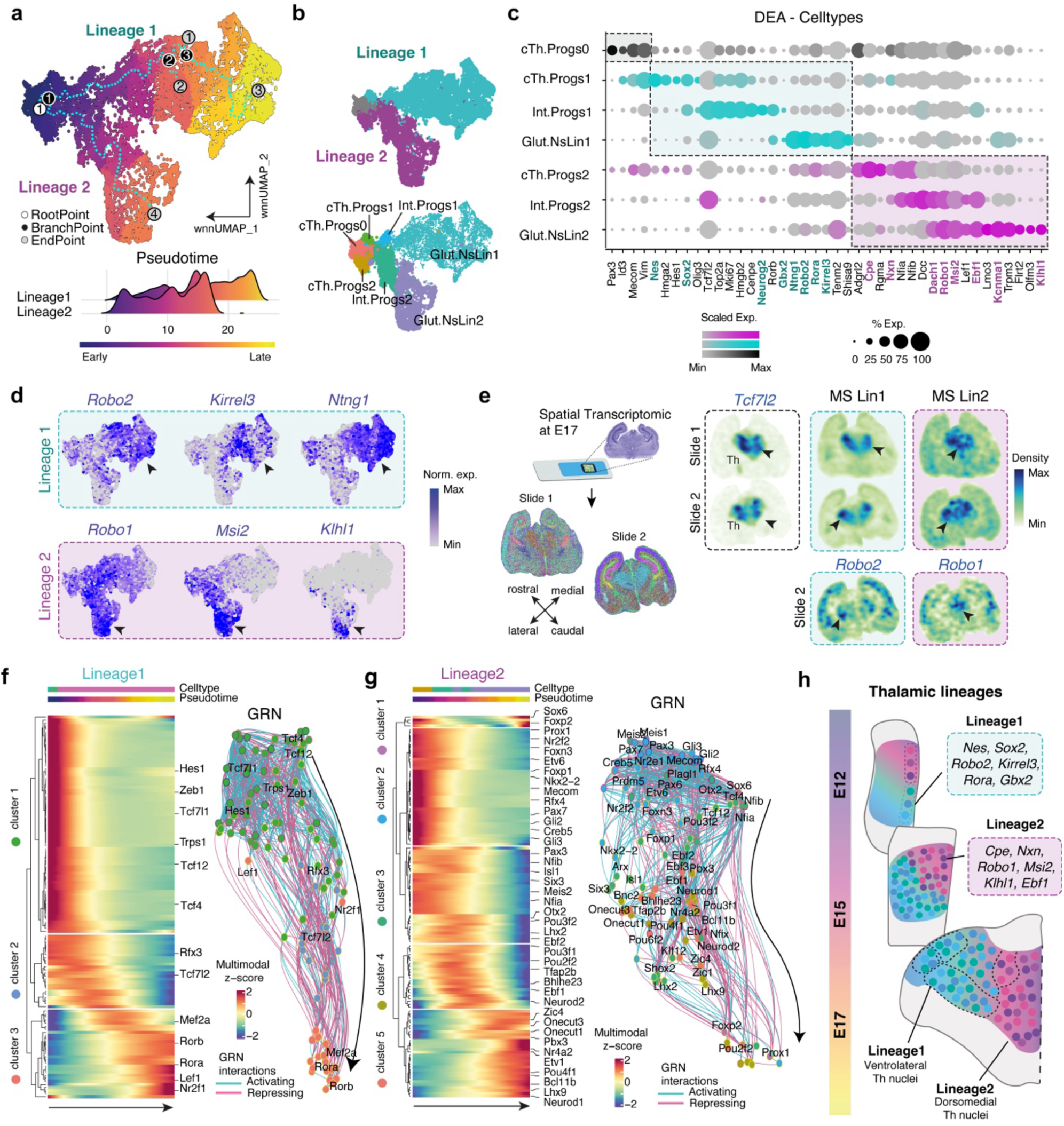
Temporally divergent developmental programmes shape modality-specific thalamic lineages. **a**, Top: UMAP visualization of inferred pseudotime trajectories, highlighting two major glutamatergic lineages. Bottom: density plot showing cell distribution along pseudotime, color-coded by lineage. **b**, Top: UMAP highlighting the two identified lineages. Bottom: lineage classification overlaid on cell type annotations. **c**, Dot plot showing expression of key marker genes across cell types within each lineage. Colored labels represent genes sets of both lineage 1 and 2. DEA: Differential Expression Analysis. **d**, Feature plots of representative genes differentially expressed between sensory/associative (Lineage 1) and motor (Lineage 2) neurons. Arrowheads indicate strong expression levels. **e**, Left: spatial transcriptomic analysis at E17 confirming anatomical segregation of the two lineages. Right: spatial gene module scores illustrating the regional distribution of lineage-specific signatures. MS: Module Score. E17: Embryonic day 17. Arrowheads indicate strong expression levels. **f**, Left: heatmap of transcription factors (TFs) in Lineage 1, clustered by expression dynamics along pseudotime. Right: UMAP embedding of the inferred TF network in Lineage 1, based on co-expression and interaction strength. Fill color indicates cluster identity; border color denotes expression-weighted pseudotime; edge color reflects predicted regulatory interactions. GRN: Gene Regulatory Network. **g**, Same as (f), for Lineage 2. **h**, Schematic summarizing the two glutamatergic thalamic lineages with specific gene markers, highlighting their developmental timing, anatomical distribution, and regulatory architecture. E: Embryonical age.

Lineage 1 was enriched for transcriptional programs associated with ventrolateral thalamic nuclei implicated in sensory processing, including both FO and HO, and expressed genes such as *Gbx2, Ntng1, Kirrel3, Rora,* and *Robo2*^27–29^. In contrast, Lineage 2 showed enrichment for dorsomedial thalamic nuclei implicated in motor and associative functions, expressing markers such as *Klhl1, Msi2, Kcnma1*, *Ebf1* and *Robo1*^27,30–32^ (Fig. 2c,d; Extended Data Fig. 5c,d). The enrichment of specific gene patterns in sensory (Lineage 1) versus non-sensory (Lineage 2) neurons was further showed by GSEA (Extended Data Fig. 6) and is consistent with previous scRNA-seq^15,33^ and clonal lineage analyses^13,34^. Whole-thalamus spatial transcriptomics at E17 further confirmed anatomical segregation, with Lineage 1 corresponding to ventrolateral domains encompassing sensory nuclei such as VPM, dLG, vMG or VPL, and Lineage 2 to dorsomedial territories including motor nuclei such as VA, VL or VM, and associative nuclei such as MD and CM (Fig. 2e; Extended Data Fig. 7). Together, these data demonstrates that sensory and non-sensory thalamic domains emerge from spatially and molecularly distinct lineages.

To elucidate the gene regulatory logic underlying these divergent fates, we reconstructed lineage-specific gene regulatory networks (GRNs) by integrating chromatin accessibility and gene expression through transcription factor (TF)–peak interaction models^35^. This analysis revealed distinct TF modules driving lineage specification, visualized via heatmaps and UMAP embeddings of TF activity across pseudotime (Fig. 2f,g). Sensory Lineage 1 progression was driven by TFs including *Hes1*, *Mef2a*, *Rora*, or *Rorb*, whereas non-sensory Lineage 2 was regulated by neurogenic TFs such as *Onecut1* and *Onecut2,* as well as by *Nr2f2*, *Foxp2*, *Ebf1* or *Neurod1*. Notably, the TF activity profile of Lineage 2 was higher than that of Lineage 1, suggesting it represents a population still undergoing active specification rather than one that is fully differentiated.

To probe causal regulators of fate decisions, we performed *in silico* TF perturbation analysis^36^. Simulated knockouts identified *Hes1* and *Neurod1* as key drivers of Lineage 1 and Lineage 2, respectively (Extended Data Fig 8a). Disruption of these TFs reversed developmental velocity within their respective branches (Extended Data Fig. 8b,c), indicating their essential and mutually exclusive roles in lineage progression. Downstream network analysis revealed that *Hes1* regulates either positively or negatively *Rora, Rorb* and *Mef2a* in Lineage 1, whereas *Neurod1* targets *Onecut1, Foxp2*, and *Ebf1* in Lineage 2 (Extended Data Fig. 8d,e). Together, these findings define the molecular programs and regulatory hierarchies that guide the early divergence of sensory and motor/associative thalamic circuits from different glutamatergic lineages (Fig. 2h).

### First-order sensory nuclei emerge from a high-order default state

Thalamic sensory neurons are functionally categorized into FO and HO nuclei, depending on whether their driver input arises from peripheral input (FO) or cortical feedback (HO). Prior studies suggest that HO identity emerges earlier than FO during development, raising the possibility that HO neurons represent a default or precursor state in the sensory lineage^16,29^. To test this hypothesis, we combined single-cell multiomic profiling with *in vivo* clonal fate mapping to reconstruct lineages relationships and developmental trajectories.

Within sensory Lineage 1, trajectory analysis of transcriptomic and chromatin accessibility profiles revealed a bifurcating developmental branch extending from shared progenitors toward two transcriptionally specific terminal states (Fig. 3a,b; Extended Data Fig. 9a). To determine whether FO and HO neurons arise from a common lineage, we performed barcoded lineage tracing at E11 using TrackerSeq^37^, which simultaneously captures transcriptomes and clonal relationships. We targeted barcoding to apical progenitors lining the third ventricle and isolated nuclei from the caudal thalamus at postnatal day (P)2 for snRNA-seq (Fig. 3c–e). We profiled 9,794 high-quality cells, which were primarily glutamatergic neurons (Extended Data Fig. 9b–e), and annotated them as FO or HO based on established marker genes^16,29^ (Extended Data Fig. 9f–i).

**Fig. 3.**
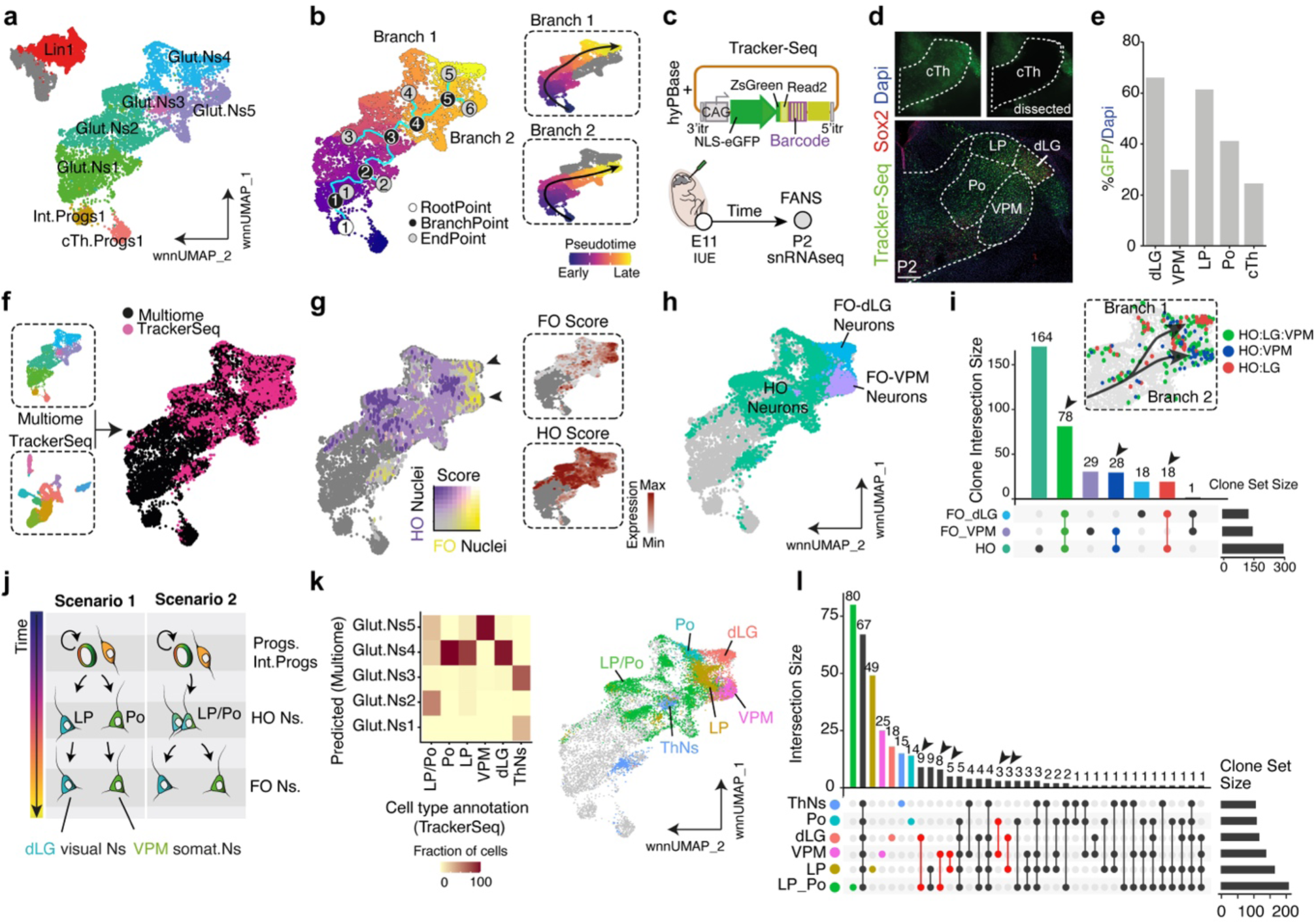
Developmental trajectories of sensory thalamic nuclei emerge from a higher-order default state. **a**, UMAP highlighting isolation of Lineage 1, corresponding to the sensory/associative thalamic lineage. **b**, Left, Pseudotime trajectory analysis reveals bifurcation into two developmental branches corresponding to distinct sensory identities. Right, UMAP showing the trajectory of every branch isolated. **c**, Top: schematic of the TrackerSeq plasmid design. Bottom: experimental workflow showing *in utero* electroporation at E11 and tissue collection at P2 for single-nucleus RNA-seq (snRNA-seq) after nuclei sorting (FANS). **d**, Top: coronal sections showing expression of TrackerSeq construct (GFP) in the caudal thalamus. Bottom: P2 coronal section showing postnatal distribution of labeled cells. **e**, Quantification of GFP/DAPI fluorescence intensity across thalamic subregions. **f**, UMAP showing label transfer from the TrackerSeq dataset to the multiome thalamic reference. **g**, Feature plot showing FO and HO nucleus assignments based on gene module scoring. **h**, UMAP embedding with final cluster annotations in the multiome dataset, incorporating lineage tracing data. **i**, UpSet plot displaying clonal relationships across cell clusters: top panel shows clone combinations; right panel shows total clone counts per cluster. Inset: UMAP illustration of clones with sibling cells traversing mixed developmental trajectories annotated with FO and HO labels. **j**, Schematic summarizing possible lineage differentiation scenarios inferred from the clonal data. **k**, Left: cross-comparison of thalamic nuclei identity predictions between TrackerSeq and the multiome dataset. Right: feature plot visualizing TrackerSeq-labeled cell types projected onto the multiome UMAP. **l**, UpSet plot as in (i), showing clone combinations and frequencies across predicted thalamic identities. Scale bar, d 500 μm.

Integration of TrackerSeq (P2) and developmental multiomic datasets (E12, E15, and E17) positioned barcoded HO (e.g. *Pde4b, Plcb1, Vwc2l, Kcnc2*)^16,29^ and FO (e.g. *Slc18a2, Kcnd3, Pde7b, Camk4*)^16,29^ neurons along the two branches of the Lineage 1 trajectory (Fig. 3f,g; Extended data Fig 9i). HO neurons retained an immature transcriptional profile, whereas FO neurons, including those from dLG and VPM, localized to the distal ends of the bifurcated branches. Notably, along the trajectory, dLG and VPM neurons acquired transcriptionally distinct identities (Fig. 3f–h; Extended data Fig 9j), supporting the model in which FO neurons emerge from a shared, earlier HO-like state.

Clonal mapping across the multiomic dataset revealed that most clones remained within a single transcriptional identity (Extended Data Fig. 9k). However, early barcoding frequently yielded mixed clones spanning multiple cell state, including both HO and FO fates. Notably, we identified strong clonal coupling between HO and FO populations: 78 clones included both HO and FO (VPM and dLG), 28 clones spanned HO and FO/VPM, and 18 clones included HO and FO/dLG (Fig. 3i). Binomial test did not reveal a significant deviation from an equal probability distribution (*P*=0.184), suggesting that HO neurons have a similar likelihood of adopting either sensory FO fate. Strikingly, only one clone contained both FO types (VPM and dLG) without an HO contribution, indicating that HO neurons are a necessary intermediate in the generation of FO neurons.

We next asked whether HO identity represents a fate-restricted precursor to specific FO subtypes (scenario 1), or whether they retain multipotency for both sensory modalities (scenario 2) (Fig3j). Early in the Lineage 1 trajectory, immature neurons showed a mixed HO identity, sharing lateral posterior (LP) and posteromedial (Po) markers, suggesting a pre-specified intermediate state. Later in the lineage annotation distinguished LP or Po based on subtype-specific markers (Fig. 3k; Extended Data Fig. 9g,h). Clonal analysis revealed that the pre-specified mixed HO identity give rise to LP, Po, dLG and VPM (Fig. 3l). Remarkably, defined LP and Po neurons also retain this capacity and could give rise to either dLG or VPM fates with equal probability (Fig. 3l; *P*=0.491, Fisher’s exact test), indicating that HO identity constitute a default intermediate state with multipotent capacity to generate either modality-specific FO neurons.

### Sp9 defines visual thalamic identity via lineage-specific transcriptional control

The specification of individual sensory thalamic nuclei appears to follow a hybrid developmental model, wherein HO identity retains plasticity before committing to modality-specific FO fates. To uncover the molecular mechanisms guiding this transition, we performed GRN inference on our single-cell multiomic data, mapping the transcriptional dynamics along trajectories within sensory Lineage 1.

While early progenitors exhibited broadly shared TF activity, lineage divergence was marked by progressive restriction of regulatory programs specific to each FO fate. Notably, *Sp9, Zic2,* and *Pou2f2,* were selectively enriched along the visual (dLG) trajectory, whereas *Nr2f1, Tshz1,* and *Rorb* predominated in the somatosensory (VPM) branch (Fig. 4a,b). Consistent with prior observations^38^, we found that *Sp9* expression was spatially restricted to the developing dLG at E17.5 (Fig. 4b,c), correlating with increased chromatin accessibility at its promoter, which remained inaccessible in VPM neurons. Conversely, *Rorb* was highly express in VPM neurons and showed greater chromatin accessibility at its promoter region (Fig. 4b).

**Fig. 4.**
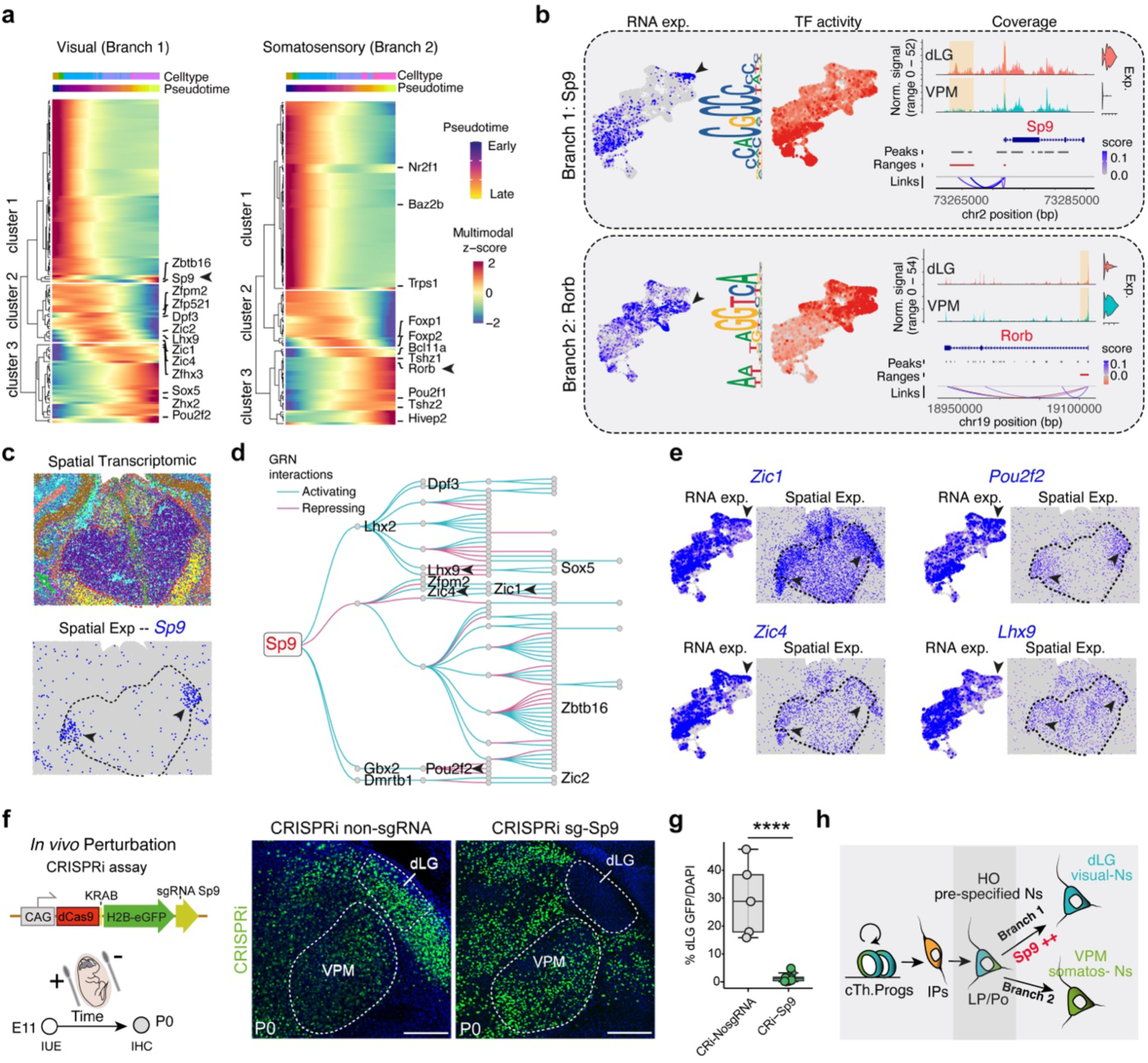
*Sp9* is required for the acquisition of visual thalamic identity. **a**, Heatmaps showing transcription factor (TF) expression dynamics along the pseudotime trajectory in visual (left) and somatosensory (right) lineages, highlighting distinct regulatory modules. **b**, Feature plots of representative TFs differentially expressed between lineages. Left, RNA expression; middle, ChromVAR-inferred gene activity; right, coverage plots showing chromatin accessibility. **c**, Top: spatial transcriptomic sub-clustering of the caudal thalamus at E17. Bottom: spatial expression of *Sp9*. **d**, Subgraph of the *Sp9*-centered gene regulatory network (GRN) showing first- and second-order transcriptional targets. Nodes represent target genes; edges are colored by inferred regulatory interaction strength. **e**, Spatial feature plots showing expression of representative *Sp9* target genes identified in the GRN. Arrowheads indicate strong expression levels. **f**, Left: schematic of CRISPR interference (CRISPRi) assay targeting *Sp9*. Right: coronal sections of the caudal thalamus at P0 showing expression following electroporation of control (CRISPRi-No sgRNA, left) or *Sp9*-targeting construct (CRISPRi-sgSp9, right). **g**, Quantification of GFP/DAPI fluorescence intensity in the dLG nucleus, showing reduced labeling upon *Sp9* knockdown (*n*= 5 CRISPRi-No sgRNA, *n*= 11 CRISPRi-sgSp9). Boxplot show the medians with the interquartile range (box) and range (whiskers). Significance was calculated using unpaired two-tailed Student T-test. ****p<0.0001. **h**, Schematic summary of findings, illustrating the lineage-specific role of *Sp9* in specifying visual thalamic identity. Scale bar, f 250 μm.

Interrogation of the dLG-specific GRN revealed that *Sp9* regulates a cascade of transcriptional targets (either activating or repressing) associated with visual thalamic identity^28^, including *Gbx2*, *Zic1, Zic4, Pou2f2* and *Lhx9* (Fig. 4d,e, Extended Data Fig. 10a,b). To assess functional relevance of *Sp9*, we performed *in silico* perturbation analysis, which predicted a lineage-specific requirement for *Sp9* in dLG specification (Extended Data Fig. 10c,d). We experimentally validated this prediction by targeting *Sp9* using CRISPR interference (CRISPRi) which was *in vivo* electroporated in the thalamus at E11 (Fig. 4f). Analysis at P0 showed that GFP-positive neurons, and therefore knockdown for *Sp9*, failed to populate the dLG nucleus (Fig. 4f and g), indicating that *Sp9* downregulation disrupts the acquisition of dLG-specific transcriptional programs. Furthermore, CRISPRi-mediated downregulation of *Sp9* in dissociated thalamic neurons led to upregulation of higher-order (HO) markers, such as *Vwc2l* and *Plcb1*, compared to controls (Extended Data Fig. 10e). Together, these results identify *Sp9* as a lineage-restricted determinant of visual thalamic identity.

## Discussion

Our study reveals that thalamic sensory and non-sensory (motor/associative) circuits arise from two early-diverging, lineage-restricted trajectories, each governed by distinct transcriptional programs and temporal dynamics. By integrating single-cell multiomics with barcoded lineage tracing, we demonstrate that, within the sensory branch, both FO somatosensory and visual neurons originate from a common progenitor pool. These progenitors are primed toward HO identity before differentiating into distinct subclasses of FO neurons, challenging classical models that proposed a largely homogeneous progenitor pool undergoing gradual diversification^11,13^.

While prior single-cell studies captured regional thalamic identities both in mice and humans^15,17,29^, they did not fully resolve the lineage architecture or regulatory logic underlying thalamic circuit assembly. Here, we show that thalamic progenitors are spatially pre-patterned along ventrolateral (sensory) and dorsomedial (motor/associative) axes. Sensory nuclei, including dLG, VPM, LP, and Po, emerge from progenitors with chromatin signatures associated to TFs such as *Hes1*, *Gbx2*, *Sox2 a*nd *Rora*, whereas motor nuclei (e.g., VL, VA, VM) and associative nuclei (e.g., CM, MD) arise from different lineages regulated by *Nr2f2*, *Lhx2*, *Ebf1* and *Neurod1*. These findings support a model in which intrinsic lineage programs, rather than postmitotic stochastic fate decisions or solely extrinsic cues^14,34^, guide functionally specialized thalamic development.

The high resolution multiomic approach presented here reveals two transcriptionally and spatially divergent lineages within the caudal glutamatergic thalamic domain (Prog2), each giving rise to functionally segregated thalamic nuclei. We identify a sensory lineage (cTh.Prog1), marked by *Sox2*, *Robo2*, *Gbx2*, *Kirrel3*, and *Nes*, which generates both high-order (LP/Po) and first-order (dLG, VPM) sensory nuclei. In contrast, a non-sensory lineage (cTh.Prog2), enriched for *Msi2*, *Robo1*, *Klhl1*, *Nxn*, and *Cpe*, give rise predominantly to motor and associative nuclei, including VL/VA, MD and CM.

Our findings parallel those reported by Govek et al^15^, who, based on single-cell transcriptomic profiling, described two major progenitor populations in the caudal thalamus that map onto overlapping sets of mature thalamic nuclei. While both studies converge on the existence of dual progenitor domains with overlapping transcriptional features, key differences emerge. Notably, in Govek et al.’s model, higher-order and first-order sensory nuclei appear to derive from separate progenitor pools. In contrast, our lineage-resolved data demonstrate that all sensory thalamic nuclei— regardless of their FO or HO classification—originate from the same caudal sensory progenitor domain. Consistent with this, recent work in the developing human thalamus supports a model in which FO and HO sensory thalamic neurons arise from a shared progenitor pool and progressively acquire distinct molecular identities^17^. This cross-species convergence reinforces the evolutionary conservation of the lineage architecture underlying sensory thalamic nucleogenesis.

Our data show that non-sensory thalamic nuclei arise from a distinct dorsomedial progenitor pool, clearly separated from the sensory lineage. We also uncover temporal asymmetry between lineages: sensory neurons initiate differentiation earlier, while non-sensory neurons remain in an immature transcriptional state during embryogenesis. Within the sensory lineage, HO neurons emerge first and serve as a developmental default state from which FO neurons later differentiate. This hierarchical relationship was supported by pseudotime analysis and *in vivo* lineage tracing and aligns with prior findings that FO identity acquisition is critically influenced by peripheral input^29^. Notably, this developmental hierarchy is mirrored in the adult thalamus, where FO and HO identities showed transcriptional profiles that are more similar at order level than across sensory-modality^10^. Furthermore, our data further identify discrete developmental trajectories from HO neurons to visual (dLG) and somatosensory (VPM) FO nuclei. These branches are transcriptionally defined and clonally validated, indicating that the transition from HO to FO involves fate-restrictive events that are tightly controlled.

Building on previous observations^38^, we show that *Sp9* functions as a lineage-specific regulator of visual thalamic identity. While *Sp9* has been implicated in specifying other neuronal subtypes^39,40^, we now demonstrate its selective requirement for dLG fate, without affecting VPM development. *Sp9* activates a transcriptional axis, including *Pou2f2*, *Zic1*, and *Zic4,* that defines the GRN guiding visual thalamic specification.

These insights have broader implications for understanding the developmental origins of thalamocortical dysfunction in neurodevelopmental disorders. Disruptions in FO thalamic circuits are associated with impaired sensory processing in autism spectrum disorder, while alterations in HO thalamus are linked to associative deficits in schizophrenia^2^. The lineage-specific transcription factors and regulatory modules we identify offer a blueprint for interrogating disease mechanisms and may inform future strategies for circuit-level reprogramming.

Beyond their clinical relevance, the spatial pre-patterning of thalamic progenitors into ventrolateral (sensory) and dorsomedial (motor/associative) domains, coupled with distinct temporal dynamics, suggests that thalamic diversity arises through early, lineage-restricted programs. This organizational principle provides a mechanistic framework for understanding how discrete thalamic outputs are developmentally matched to cortical targets with specialized functional roles.

In summary, we define the transcriptional and lineage logic that orchestrates thalamic circuit specification. This framework not only refines our understanding of sensory brain development but also provides molecular entry points for probing and manipulating thalamocortical function in health and disease.

## Methods

### Mouse strains

All animal procedures were approved by the Animal Welfare Committee of the University Miguel Hernández and carried out in accordance with Spanish and European Union guidelines for the care and use of laboratory animals. Mice were housed under specific pathogen-free conditions on a 12-h light/dark cycle, with ad libitum access to food and water. No sex-dependent differences were observed, and both male and female embryos were used interchangeably across experiments. The number of animals used per experiment is provided in the corresponding figure legends. All transgenic lines were maintained on either ICR/CD-1 or C57BL/6J backgrounds and genotyped by PCR. The morning of vaginal plug detection was designated embryonic day 0.5 (E0.5).

### *In utero* FlashTag injection

*In utero* labeling was performed using FlashTag as previously described^18^. Pregnant mice at embryonic day 11 (E11) were anesthetized with 2–3% isoflurane and placed on a 37 °C heated surgical platform. Following laparotomy, the uterine horns were exposed, and 414–483 nL of 6 mM FlashTag (carboxyfluorescein succinimidyl ester; CellTrace CFSE, Life Technologies, C34554) was injected into the lateral ventricles using a nanoinjector fitted with pulled, beveled, and dust-free glass micropipettes. After injection, the uterine horns were returned to the abdominal cavity, and the peritoneal wall and skin were sutured in separate layers. Mice were monitored postoperatively and placed on a warming pad until full recovery from anesthesia.

### *In utero* electroporation

*In utero* electroporation was performed as previously described^41^. Pregnant females at E11.5 were deeply anesthetized with isoflurane, and a midline laparotomy was conducted to expose the uterine horns. Embryonic brains were visualized through the uterine wall using an optic fiber light source. Approximately 414–483 nL of plasmid DNA (diluted in endotoxin-free TE buffer containing 0.002% Fast Green FCF; Sigma) was injected into the lateral ventricle of each embryo. Electroporation was carried out by positioning circular tweezer electrodes (5 mm diameter, Sonidel) across the uterine wall and delivering five square-wave pulses (50 V, 50 ms duration, 1 Hz) using a pulse generator (Nepa Gene, Sonidel). Embryos were returned to the abdominal cavity, and dams were sutured and monitored during recovery.

### Histology and image acquisition

For embryonic tissue, dissected brains were fixed overnight in 4% paraformaldehyde (PFA) at 4 °C. For postnatal stages, mice were transcardially perfused with 0.01 M phosphate-buffered saline (PBS) followed by 4% PFA. Brains were embedded in 3% agarose (in 0.01 M PBS) and sectioned at 80 µm thickness using a vibratome (VT1200S, Leica). Sections were blocked for 1 h at room temperature in a solution containing 1% bovine serum albumin (BSA), 2% donkey serum, 2% goat serum, and 0.2% Triton X-100 in 0.01 M PBS. Primary antibody incubation was performed overnight at 4 °C, followed by 2 h incubation with secondary antibodies at room temperature. Nuclei were counterstained with fluorescent nuclear dye 4′,6-diamidino-2-phenylindole (DAPI) (Sigma-Aldrich). Images were acquired with a Leica K5 camera mounted on a Leica DMi8 microscope with large volume computational clearing post processing and quantified using ImageJ. High resolution images were acquired with a Zeiss LSM 880-Airyscan Elyra PS.1 inverted super-resolution confocal microscope, in Airyscan mode, with standard Airyscan deconvolution postprocessing.

### Tissue preparation and cellular dissociation for single-cell multiome sequencing

Embryonic thalamic tissues were microdissected from four embryos per timepoint (E11.5, E14.5, and E16.5) and pooled for single-cell Multiome-seq. Tissue was dissected in cold dissociation medium containing 20 mM glucose, 0.8 mM kynurenic acid, 0.05 mM d,l-2-amino-5-phosphonovaleric acid (APV), 50 U/ml penicillin, 0.05 mg/ml streptomycin, 0.09 M Na₂SO₄, 0.03 M K₂SO₄, and 0.014 M MgCl₂. Samples were enzymatically digested in dissociation medium supplemented with 0.16 mg/ml l-cysteine and 70 U papain (Sigma-Aldrich), pH 7.35, at 37 °C for 30 min with repeated shaking.

Enzymatic activity was halted with ovomucoid (0.1 mg/ml) and BSA (0.1 mg/ml) in dissociation medium (pH 7.35, room temperature). Tissues were washed in ice-cold Opti-MEM (Life Technologies) containing 20 mM glucose, 0.4 mM kynurenic acid, and 0.025 mM APV, then mechanically triturated to generate a single-cell suspension. Cells were pelleted by centrifugation at 850 rpm for 5 min and filtered through a 35 µm cell strainer (BD Falcon). Genetically labeled cells were sorted based on green fluorescence using a FACSAria III cell sorter (BD Biosciences). Sorted cells were used for single-nucleus isolation for downstream multiomic processing.

### Single-nucleus isolation and single-cell multiome library preparation

Single-nucleus isolation was performed following the 10x Genomics Demonstrated Protocol for Single Cell Multiome ATAC + Gene Expression (CG000365). FACS-purified cell suspensions were resuspended in 800 μL of PBS without Ca²⁺/Mg²⁺, supplemented with 0.04% BSA and 1 U/μL RNase inhibitor. After centrifugation at 600 g for 7 min at 4 °C, cell pellets were lysed on ice for 5 min in 100 μL of chilled 0.1× lysis buffer [10 mM Tris-HCl pH 7.4, 10 mM NaCl, 3 mM MgCl₂, 1% BSA, 1 mM DTT, 1 U/μL RNase inhibitor, 0.1% Tween-20, 0.1% IGEPAL CA-630, and 0.01% Digitonin]. Lysis was stopped by adding 500 μL of wash buffer [10 mM Tris-HCl pH 7.4, 10 mM NaCl, 3 mM MgCl2, 1% BSA, 1 mM DTT, 1 U/μL RNase inhibitor, and 0.1% Tween-20].

Nuclei were collected by centrifugation at 500 g for 5 min at 4 °C, rinsed three times in 200 μL wash buffer, and resuspended in Nuclei Resuspension Buffer (10x Genomics) supplemented with 1 mM DTT and 1 U/μL RNase inhibitor. Filtered nuclei (35 μm strainer) were inspected microscopically to confirm single-nucleus dissociation.

Single-nucleus libraries were generated using the Chromium Next GEM Single Cell Multiome ATAC + Gene Expression platform (10x Genomics; CG000338). Nuclei were incubated with a transposase to fragment accessible chromatin and tag DNA ends with adapters. Transposed nuclei were loaded onto Chromium Next GEM Chip J (PN-1000234) with partitioning oil and barcoded gel beads. Following barcoding and amplification, ATAC and gene expression libraries were constructed separately. Library quality was assessed using the High Sensitivity DNA Kit (#5067-4626, Agilent) on a Bioanalyzer 2100 (Agilent Technologies).

Sequencing was performed at the Novogene core facility (Munich, Germany) on an Illumina NovaSeq X system. RNA-seq libraries were sequenced using a 28–10–10–90 bp read configuration. ATAC-seq libraries were sequenced with either a 50–8–16–50 bp or paired-end 2×150 bp configuration.

### Single-cell multiome data processing

Raw sequencing data were processed using CellRanger ARC (v2.0.1, 10x Genomics). RNA and ATAC reads were aligned to the GRCm38 mouse reference genome using the “cellranger-arc count” command.

Downstream analysis was performed in R (v4.2.2) using the Seurat (v4.4.1) and Signac (v1.3.0) packages. For RNA, cells were filtered using the following thresholds: nFeature_RNA < 7,500; 200 < nCount_RNA < 100,000; percent.mt < 5%. For ATAC, cells were filtered based on: 200 < nCount_ATAC < 100,000; nucleosome_signal < 2; TSS.enrichment > 1 (Extended Data Fig 1). After filtering, the dataset included 8,009 nuclei (E11.5), 8,460 nuclei (E14.5), and 9,008 nuclei (E16.5). Gene expression counts were normalized using SCTransform. Dimensionality reduction was performed via PCA on the top 3,000 highly variable genes, and uniform manifold approximation and projection (UMAP) embeddings were computed using the first 30 principal components. Doublets were identified using a consensus approach with the scDblFinder (v1.12) and scds (v1.14) packages and excluded from downstream analysis.

For ATAC data, peak calling was performed using MACS2 via the CallPeaks function in Signac. Peaks overlapping genomic blacklist regions were removed. The peak count matrix was normalized using Latent Semantic Indexing (LSI), including term frequency-inverse document frequency (TF-IDF) and singular value decomposition (SVD). The first LSI component, which correlated with sequencing depth, was excluded. Chromatin accessibility–based gene activity was quantified using the ChromVAR function (v1.20.2), aggregating accessibility across gene bodies and promoters.

### Bi-modal integrative analysis of RNA-ATAC scMultiome data

To integrate the RNA and ATAC modalities from multiomic data, we employed the weighted nearest neighbour (WNN) approach implemented in Seurat^42^. This method constructs a k-nearest neighbour graph that balances information from both modalities using weights determined for each cell. FindMultiModalNeighbors function was applied in Seurat with the integrated embeddings of RNA and ATAC as inputs. The resulting neighbour graph was used for UMAP embedding and clustering, enabling a comprehensive analysis of cellular heterogeneity and transcriptional regulation.

Cell clustering was carried out with FindNeighbors and FindClusters (resolution =0.3). Cell clusters were annotated using canonical markers for glutamatergic and gabaergic neurons, intermediate progenitors, progenitors, glioblast, pericytes, ependymals andoligodendrocyte precursor cells (OPCs).

To identify marker genes distinguishing clusters, we performed differential expression analysis using Seurat’s FindAllMarkers function on the RNA assay, applying a log fold change threshold of 0.25 and a minimum fraction of expressing cells of 0.1, with p-values adjusted using the Bonferroni correction.

### Peak–gene linkage analysis

Peak–gene associations were inferred from chromatin accessibility data using the LinkPeaks function in the Signac package, based on the strategy previously described^43^. Pearson correlation coefficients were calculated between gene expression and accessibility of nearby peaks, restricted to regions within 500 kb of the gene transcription start site (TSS). Covariates including GC content, overall peak accessibility, and peak width were included to control for technical biases. P-values were adjusted using the Benjamini–Hochberg method. Only peak–gene pairs with an adjusted P-value < 0.05 and positive correlation coefficients were retained for downstream analysis.

### DNA motif enrichment analysis

To identify putative transcription factor (TF) binding sites, motif enrichment analysis was performed on sets of differentially accessible chromatin regions. Position weight matrices (PWMs) were obtained from the JASPAR 2022 database using the getMatrixSet function in the TFBSTools package (v1.36) (collection: “CORE”; tax_group: “vertebrates”). Enrichment was performed using the FindMotifs function in Signac with default parameters. Motif activity was quantified using ChromVAR, which computes deviation scores across nuclei by aggregating accessibility profiles at motif-matching genomic loci.

### Pseudotime trajectory analysis

Pseudotime trajectories were constructed using the Monocle3 workflow (v1.3.7). The integrated Seurat object was converted into a cell_data_set using the SeuratWrappers package (v0.3.5). Dimensionality reduction and cell clustering was based on the WNN-UMAP embedding from the Seurat object. The principal graph was learned with learn_graph (use_partition = FALSE; minimal_branch_len = 15, ncenter=500), and pseudotime was assigned using the order_cells function with default settings. For branch-specific ordering, the choose_graph_segments function was used to connect the earliest and most advanced developmental states along each lineage. Only the dominant trajectory per lineage was visualized in downstream analyses. A slingshot pseudotime analysis was used in the isolated lineage 1 dataset to infer the developmental bifurcation and select the cells belonging to each branch using the Slingshot package (v2.7.0).

### RNA velocity analysis

The post-sorted bam files from CellRanger-arc output were passed to velocyto (v0.17.15) to estimate the RNA velocities of single cells. The generated loom file contained data matrices of spliced and unspliced reads and was further processed by scVelo (v0.3.2).

Seurat-processed gene expression count matrix and UMAP coordinates were converted to “AnnData” object and merged with the velocyto-derived object using scVelo.utils.merge function. The merged dataset was filtered using the scVelo.pp.filter_and_normalize with default parameters (min_shared_counts = 10, n_top_genes = 2000) and the moments were computed using scVelo.pp.moments. The velocity was then calculated using scVelo.tl.velocity (mode=stochastic). The estimated velocity vectors were projected and visualized in previously calculated embedding. The initial and terminal state likelihood based on RNA velocity information was estimated using cellrank.tl.terminal_states and cellrank.tl.initial_states functions with default parameters (weight_connectivities=0.3).

### Gene ontology over representation analysis and gene set enrichment analysis

Gene ontology (GO) over-representation analysis was performed for biological processes using the enrichGO function from the clusterProfiler package (v4.6.2), with the following parameters: pAdjustMethod = “BH”, qvalueCutoff = 0.001, minGSSize = 10, maxGSSize = 500. Gene set enrichment analysis (GSEA) was performed using clusterProfiler with the following parameters: pAdjustMethod = “BH”, pvalueCutoff = 0.2, minGSSize = 1, maxGSSize = 1000, nPermSimple=10000. The reference gene set used to perform the analysis was extracted from an available dataset^15^.

### Gene regulatory network (GRN) inference and visualization

GRNs were inferred using Pando (v1.1.1), integrating RNA and ATAC profiles from metacells as previously described^35^. Conserved peaks overlapping PhastCons elements were lifted to mm10 with liftOver (v1.18.0), and motifs from JASPAR 2022 and CIS-BP were scanned using find_motifs function. Only TFs associated with the top 2,000 variable genes were retained. Peaks were assigned to genes if overlapping the gene body or within 100 kb upstream of the transcription start site, and TF–target relationships were inferred with infer_grn. Regulatory modules were defined using find_modules function, applying ANOVA and Benjamini–Hochberg FDR correction (FDR < 0.05). GRNs were visualized with get_network_graph and plot_network_graph (UMAP layout, weighted method), where node sizes reflect PageRank centrality. Lineage-specific TF contributions were assessed by calculating absorption-probability–weighted expression scores, combining lineage-specific absorption probabilities with TF expression.

### *In silico* TF perturbation analysis

We performed TF perturbation simulations using CellOracle (v0.17.2)^36^ following the recommended pipeline (https://morris-lab.github.io/CellOracle.documentation/). For computational efficiency, 3,000 cells and 3,000 highly variable genes were subsampled from the scRNA-seq data. Preprocessing and clustering were carried out using Scanpy (v1.9.6). Diffusion maps, PAGA, and force-directed graphs were computed using default parameters. For co-accessibility network inference, scATAC-seq data were processed with Cicero (v1.16.2) using a 500 kb window. Peaks with substantial co-accessibility scores (co-accessibility ≥ [0.3]) were retained. TF–target relationships were filtered using a significance threshold of *P* < 0.001, absolute regression coefficient (“coef_abs”) as a weighting metric, and a maximum of 2,000 TF–target links. Simulated TF perturbations were modeled by setting TF expression to zero. Transition probabilities and cell state shifts were computed with CellOracle’s default settings (n_neighbors = 200, knn_random = TRUE, sigma_corr = 0.05). Developmental flow was reconstructed using diffusion pseudotime, and the perturbation effect was quantified by computing the inner product between the developmental flow vector and the simulated perturbation vector.

### TrackerSeq library preparation and electroporation

TrackerSeq is a hyperactive piggyBac (hyPBase) transposon–based^44^ library that was previously developed to be compatible with the 10x single-cell transcriptomic platform^37^. It records the *in vivo* lineage history of single cells through the integration of multiple oligonucleotide sequences into the genome. Each of these individual lineage barcodes is a 37-bp-long synthetic nucleotide that consists of short random nucleotides bridged by fixed nucleotides. This design results in a library with a theoretical complexity of approximately 4.3 million lineage barcodes (16^8^) with each barcode differing from another by at least 5 bp. The barcode library was prepared as described previously^37^ with minor modifications: replacing the original GFP reporter with two distinct fluorescent reporters, a nuclear localization signal (NLS)-eGFP and ZsGreen, into the plasmid constructs to facilitate the identification and tracking nuclear-localized and cytoplasmic fluorescence detection, respectively, of transduced cells *in vivo*.

We assessed the integrity of the TrackerSeq barcode library by sequencing to a depth of approximately 13 million reads to test whether any barcode was overrepresented. A total of 5 × 10^6^ clusters of barcodes were identified, suggesting that the barcode library has a diversity that is at least in the 10^6^ range.

Following the electroporation method described above, E11 mouse embryos were electroporated with the hyPBase -transposase and the TrackerSeq library.

### Tissue preparation and cellular dissociation for TrackerSeq experiment

To collect tissue from the thalamus of electroporated animals at P2, and perform *in vivo* lineage tracing of single cells, animals were decapitated, and their brains were dissected out in RNase-free conditions to prevent RNA degradation. The brains (three brains were pooled for each sample) were collected in ice-cold artificial cerebrospinal fluid (aCSF) solution (119 mM NaCl, 5 mM KCl, 1.3 mM MgSO4, 2.4 mM CaCl2, 1 mM NaH2PO4, 26 mM Na2HCO3, and 11 mM glucose) and cut in the vibratome in 300 μm slices. The ventrolateral thalamus was rapidly microdissected under the stereo microscope and then frozen.

Single-nucleus isolation for the TrackerSeq experiment was performed based on the Demonstrated protocol for Nuclei isolation from tissue from 10xGenomics (CG000124). The frozen tissue were then suspended with 500 μL of 0.1× lysis buffer [10 mM Tris-HCl pH 7.4, 10 mM NaCl, 3 mM MgCl2, 0,2 U/μL RNase inhibitor and 0.1% IGEPAL CA-630 (i8896 Sigma-Aldrich)] for 3 min on ice, which was stopped with chilled 500 μL wash buffer [1% PBS without Ca/Mg+2 supplemented with 1% BSA and 0,2 U/μL RNase inhibitor] and the samples were pipetted 5 times using the P1000. The nuclei were then collected by centrifugation at 500× g for 5 min at 4 °C, rinsed with 200 μL wash buffer three times, and re-suspended in wash buffer. Isolated single-cell nuclei were filtered using a cell strainer (35-μm pore size) and inspected under a microscope to ensure they were successfully dissociated into single cells for subsequent sequencing. Hoechst was added to label nuclei for FANS gating. 43.2 μl of GFP+ nuclei were sorted directly into empty 1.5 ml Eppendorf for downstream processing on the 10x Genomics Chromium platform.

### scRNAseq data generation of TrackerSeq libraries

The single-nuclei RNAseq libraries were constructed by following the 10x Genomics Chromium Next GEM Single Cell 3′ Reagent Kit v3.1 (Dual Index) following the manufacturer’s protocol (document number CG000315, 10x Genomics).

Uniquely barcoded RNA transcripts were reverse-transcribed. 3′ Gene Expression libraries were generated according to the manufacturer’s user guide with use of Chromium Next GEM Single Cell 3’ Kit v3.1, 4 rxns v3.1 kit (PN-1000269), Chromium Next GEM Chip G Single Cell Kit, 16 rxns (PN-1000127) and Dual Index Kit TT Set A (PN-1000215) (10x Genomics). 3′ Gene Expression Libraries were quantified with Agilent Bioanalyzer.

The lineage/TrackerSeq barcode library amplification process followed the previously outlined procedure^37^, using the standard NEB protocol for Q5 Hot Start High-Fidelity 2X Master Mix (#M094S) in a 50-μl reaction, with 10 μl of cDNA as the template. To be specific, each PCR reaction contained the following components: 25 μl Q5 High-fidelity 2X Master Mix, 2.5 μl of a 10 μM P7_indexed reverse primer, 2.5 μl of a 10 μM i5_indexed forward primer, 10 μl of molecular-grade H20, and 10 μl of cDNA. Libraries were purified with a dual-sided selection using SPRIselect (B23318, Beckman Coulter) and quantified with Agilent Bioanalyzer.

Transcriptome and TrakerSeq barcode libraries were sequenced on an Illumina NovaSeq X platform at Novogene Co. Ltd. Genomics core facility (Munich, Germany), with 28-10-10-90 bp configuration for RNA-seq libraries.

### Single-cell RNA-seq data processing for TrackerSeq

Single-nucleus RNA-seq data were processed using CellRanger (v8.0.1, 10x Genomics). Raw FASTQ files were aligned to the GRCm38 mouse genome reference using default parameters. Filtered gene-barcode matrices were imported into Seurat (v4.4.1) in R (v4.2.2) for downstream analysis. Cells were filtered using the following thresholds: 300 < nFeature_RNA; percent.mt < 5%. Doublets were identified and excluded from downstream analysis (see above).

Normalization and scaling were performed using Seurat’s standard workflow. PCA was computed on the top 3,000 most variable genes. Dimensionality reduction was performed using UMAP on the first 30 PCs, and cell clustering was carried out with FindNeighbors and FindClusters (resolution =0.3). Cell clusters were annotated using canonical markers for neurons, astrocytes, oligodendrocyte precursor cells (OPCs), and microglia. Differential expression analysis was performed to identify marker genes distinguishing clusters (see above).

Lineage barcode FASTQ files were processed using the TrackerSeq pipeline (https://github.com/mayer-lab/TrackerSeq), as previously described^37^. Parameters were set to nReads = 10, nUMI = 2, and nHamming = 4. Barcode-extracted FASTQs were used to generate sparse matrices with cell barcodes as rows and clone IDs as columns. Biological replicates were processed separately and merged following clone ID assignment.

### *In vivo* CRISPR interference (CRISPRi)

CRISPick^45^ was used to design guide RNAs (gRNAs) targeting *Sp9* for SpyoCas9-mediated CRISPRi, based on established on-target scoring models^46^. The top three gRNA sequences were ordered as single-stranded oligonucleotides (Sigma-Aldrich) with homology arms for cloning.

Guide sequences were cloned into the pX458-CAG-dCas9-KRAB-MECP2-H2B-GFP-NogRNA backbone by BbsI digestion and NEBuilder HiFi DNA assembly, as previously described^33^. The CAG promoter was inserted into the Ef1a-dCas9-KRAB-MECP2-H2B-GFP-NogRNA plasmid (Addgene #175573) using NEBuilder HiFi.

Plasmids (1.5 μg/μL) were delivered into the embryonic thalamus at E11 via in utero electroporation (see above). Electroporations were performed using either control (CRISPRi-No gRNA) or gene-targeting (CRISPRi-sgRNA) constructs.

To validate CRISPRi knockdown, primary thalamic neurons were cultured from E14.5 embryos. 48 hours after plating, cells were transfected with CRISPRi plasmids using LipoD293 DNA in vitro Transfection Reagent (Ver. II) (SignaGen Laboratories) according to the manufactureŕs instructions. Cells were harvested for RNA extraction 48 hours after transfection.

### RNA extraction and quantitative real-time PCR

Total RNA was extracted using a Direct-zol RNA miniprep kit (Zymo Research). Then, cDNA was synthesized from 1 μg of RNA using the High-Capacity cDNA Reverse Transcription Kit (Applied Biosystems) following the manufacturer’s instructions. qPCR was performed using the StepOnePlus Real-Time PCR System (Applied Biosystems) with Power SYBR Green PCR Master Mix and gene-specific primers. All reactions were run in triplicate. Ct values were normalized to Gapdh, and relative gene expression was calculated using the ΔΔCt method.

### Spatial transcriptomics

Spatial transcriptomic profiling was performed using the Curio Seeker 10×10 Kit (Curio Biosciences, CB-000578-UG v1.3). Brains were washed in PBS with 2 U/mL RNase inhibitor, incubated in tissue-freezing medium (Electron Microscopy Sciences), embedded in BEEM flat molds, and snap-frozen on dry ice. Samples were cryosectioned at −18 °C using a Leica cryostat (CM1860 UV, Leyca

Biosystems) at 10 μm thickness. Sections were carefully mounted on the Curio Seeker 10×10 mm tile and melted by placing a finger under the tile. A 30-μm-thick section of the CryoCube was placed on top of the tissue section and melted similarly.

Two sections per brain were mounted per tile. RNA extraction and library preparation followed the manufacturer’s protocol. Tagmentation was performed using 4.8 ng cDNA and the Nextera XT Library Prep Kit (Illumina, FC-131-1024). Libraries were pooled and sequenced on a NovaSeq X Plus platform (Illumina) using paired-end 150 bp reads (150–8–8–150 bp configuration).

Spatial gene expression matrices were processed using the Curio Seeker pipeline (v3.0). Cell type assignment was performed using RCTD via the spacexr package (v2.2.1, doublet mode), using an E9.5 thalamic dataset as a reference^22^ with the multiomic E11.5 dataset data to assign cell type weights to spatial coordinates.

### Statistics and reproducibility

All high-throughput experiments were performed using at least *n* = 2 biological replicates per group. No statistical methods were used to predefine sample size. Low-quality cells were excluded from all analyses. Statistical significance was assessed using the Wilcoxon rank-sum test unless otherwise specified. Multiple comparisons were corrected using the Benjamini–Hochberg procedure where appropriate. All details regarding replicate numbers, statistical tests, and thresholds are provided in the main text and figure legends. For data that conforms to a normal distribution, an unpaired two-tailed Student’s t-test was employed to compare the two groups.

## Data availability

All data supporting the findings of this study are available within the article and its Supplementary Information. The raw and processed single-cell multiome (scRNAseq and scATACseq) data, snRNAseq data from the TrackerSeq experiment and Spatial Transcriptomic data generated in this study have been deposited in the NCBI Gene Expression Omnibus (GEO) and will be available upon publication. Additional data and materials necessary for reanalysis are available from the corresponding author upon request.

## Acknowledgements

We thank Luis Rodríguez Malmierca for technical support, and Francisco José Martini, Teresa Guillamón-Vivancos and Eduardo Leyva-Diaz for insightful comments on the manuscript. We are also grateful to members of the López-Bendito laboratory for stimulating discussions throughout the course of this work. We thank Denis Jabaudon for his supervision of Awais Javed, for generously sharing reagents, and discussions on the project. We thank Christian Mayer for generously providing the original TrackerSeq plasmids. This study was supported by grant RTI2018-102260-B-I00 from the MCIU/AEI/10.13039/501100011033 to J.L.-A.; and by the European Research Council under the European Union’s Horizon 2020 research and innovation programme (ERC-2021-ADG-101054313, SPONTSENSE), La Caixa Foundation (HR20-00023-LA CAIXA), PID2021-127112NB-I00 grant from the MCIN/AEI /10.13039/501100011033 (G.L-B.), as well as the Generalitat Valenciana, Conselleria d’Educació, Universitats i Ocupació (PROMETEO 2021/052) to G.L-B. L.P. was supported by the Spanish Ministry of Science and Innovation (PRE2019-087666). The Institute of Neurosciences is a Severo Ochoa Center of Excellence (grant CEX2021-001165-S, funded by MCIU/AEI/10.13039/501100011033).

## Author contributions

Conceptualization, L.P-A, J.L-A and G.L-B.; methodology, L.P-A, B.A-B, E.W, A.T-M and A.J.; data curation, L.P-A and E.W; writing – original draft, L.P-A, J.L-A and G.L-B.; writing – review & editing, L.P-A, E.W, A.J, J.L-A and G.L-B.; funding acquisition, G.L-B.; resources G.L-B.; supervision, J.L-A and G.L-B.

## Declaration of generative AI and AI-assisted technologies in the writing process

During the preparation of this work the authors used Chat-GPT to streamline some parts of the text. After using this tool, the authors reviewed and edited the content as needed and take full responsibility for the content of the publication.

