## Supplementary Data Figures for "Early lineage divergence segregates sensory and non-sensory thalamic circuits"

### Extended Data Figures

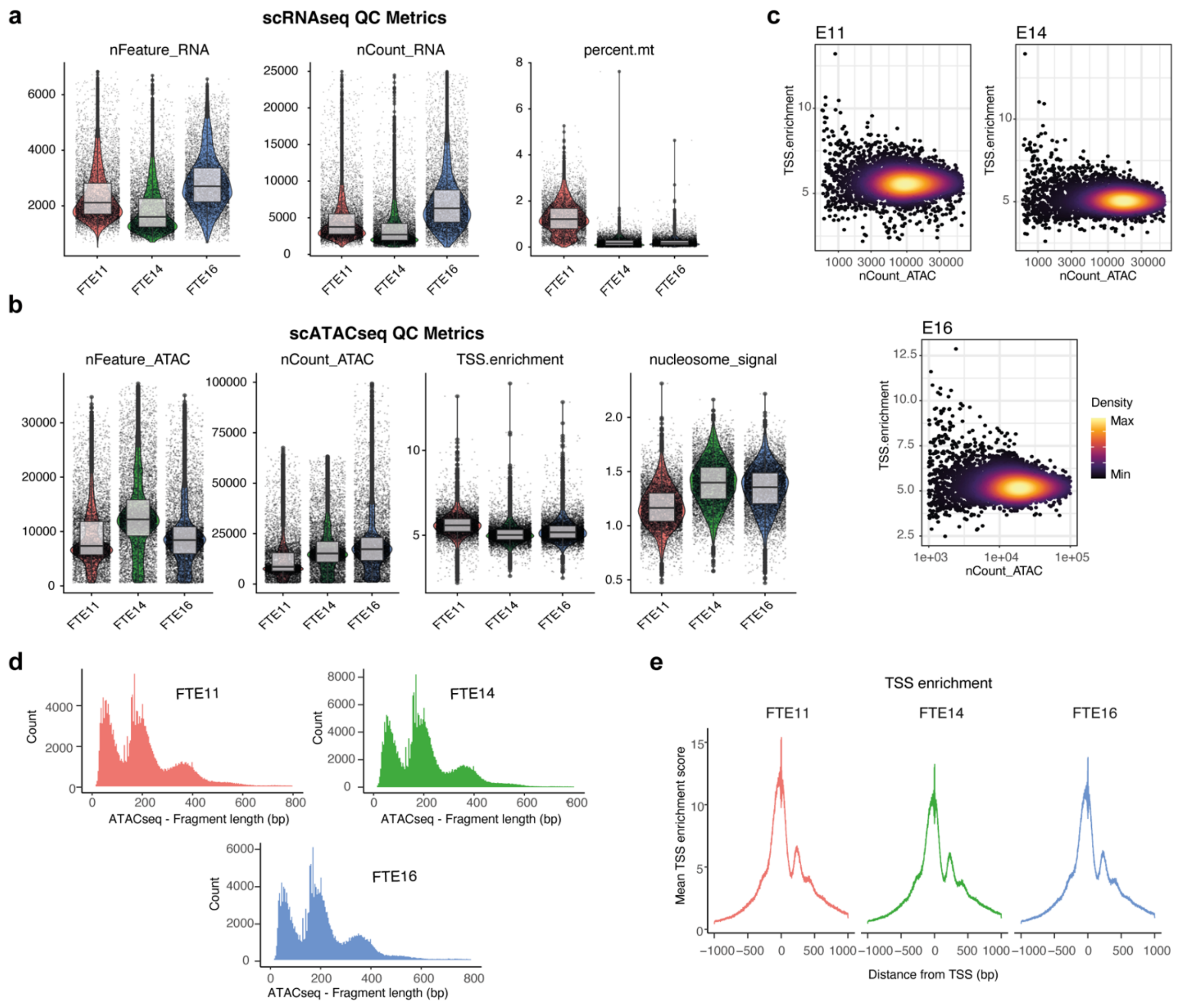


**Extended Data Fig. 1. Data quality of snATAC-seq and snRNA-seq libraries. a**,**b**, Violin plots show the distribution of several scRNA-seq (a) and scATAC-seq (b) quality metrics for each cell. scRNA-seq metrics include the total number of genes detected, total number of RNA reads, and percentage of counts from mitochondrial genes (from left to right). scATAC-seq metrics include total number of ATAC fragments, transcription start site (TSS) enrichment and nucleosome signal (from left to right). Each plot is separated by developmental stage (x-axis). E: embryonic day. **c**, Hexbin plots show density of cells stratified by total number of fragments (x-axis) and TSS enrichment (y-axis). Colour represents density. **d**, Density plot shows the distribution of fragment size for each sample. **e**, Plot shows the normalized fragment count in each positive relative to TSS (bp) for each sample. **(Related to Fig 1)**

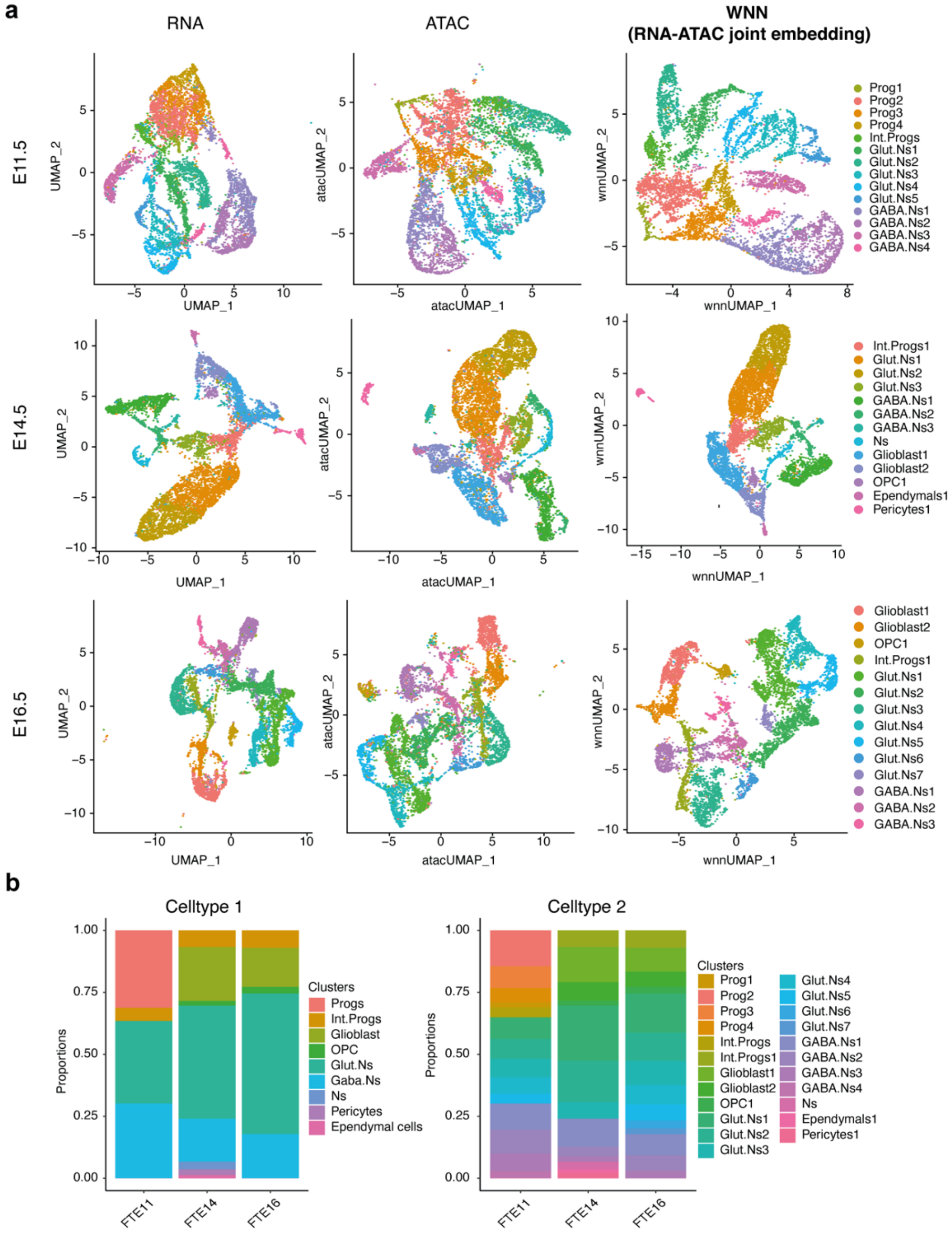


**Extended Data Fig. 2. Cell type identification of thalamic dataset at scMultiome level at three developmental stages. a**, UMAP visualization of single cells defined by scRNA-seq and scATAC-seq data and the Weighted Nearest Neighbour (WNN) analysis (RNA-ATAC joint embedding), respectively at three developmental stages. Cell type annotations are derived from the WNN umap plot. **b**, Cell type proportions in each age group. Left: a simplified manner of encompassing the cell clusters. Right: cell cluster annotation for the whole dataset together. **(Related to Fig 1)**

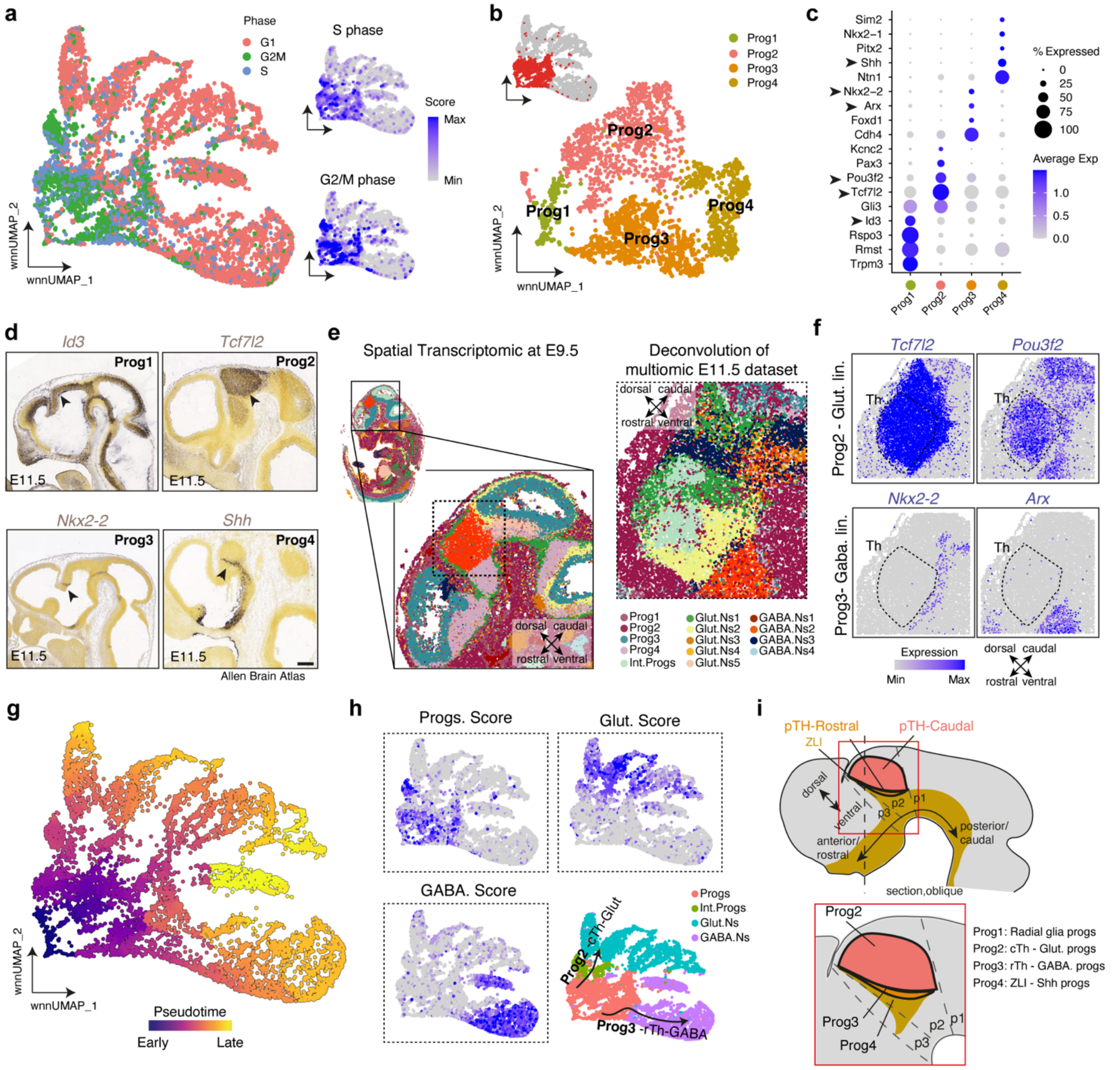


**Extended Data Fig. 3. Diverse progenitor populations give rise to divergent Glutamatergic and GABAergic thalamic lineages.** **a**, Left: UMAP visualization shows the cell cycle expression makers. Right: Feature plots show the expression of cells in S-phase and G2/M phases. **b**, UMAP visualization of progenitor population isolated from E11.5 multiomic dataset. **c**, Dotplot of normalized average expression of top genes. **d**, *In-situ* hybridization images of selected genes extracted from Allen Brain Atlas: Developing Mouse Brain. Arrowheads indicate strong expression levels. **e**, Spatial transcriptomic dataset at E9.5 of a whole embryo extracted from available dataset[22]. Dash-lined inset**,** Spatial deconvolution of cells identified in E11.5 dataset onto their spatial location. **f**, Feature plots of marker genes for Prog2 – Glut lin (top) and Prog3 – GABA lin (bottom) at spatial level. **g**, UMAP visualization showing the pseudotime analysis. **h**, Left and top-right, Feature plots showing the Gene-set module score of cell type markers (Progs Score: *Mki67*, *Top2a*, *Cenpe* and *Hmga2*; Glut. Score: *Slc17a6, Ebf1* and *Ebf3*; GABA Score: *Nrxn3*, *Gad1*, *Gad2* and *Dlx1*). Bottom-right, Schematic illustration of glutamatergic and GABAergic trajectories inferred. **i**, Schematic illustration of results. Scale bars, d 500 µm. **(Related to Fig 1)**

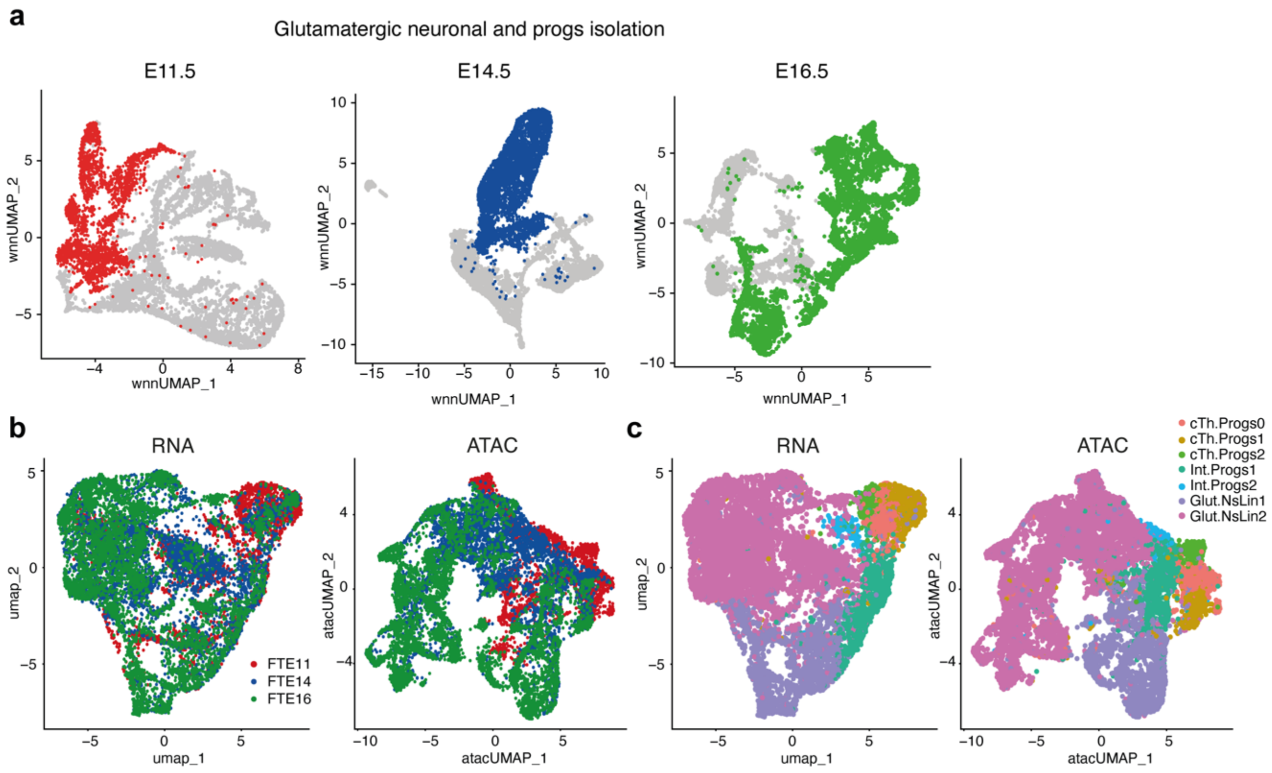


**Extended Data Fig. 4. Glutamatergic neuronal isolation and bi-modal integrative analysis of RNA-ATAC scMultiome data. a**, UMAP visualization of Weighted Nearest Neighbour (WNN) analysis at three timepoints, highlighting the isolation of glutamatergic and progenitor cell population. **b**, UMAP displaying the three isolated time points in the integrated scRNA-seq (left) and scATAC-seq (right) data. **c**, UMAP visualization based on integrated scRNA-seq (left) and scATAC-seq (right) data represent cells coloured by each cell type. **(Related to Fig 1)**

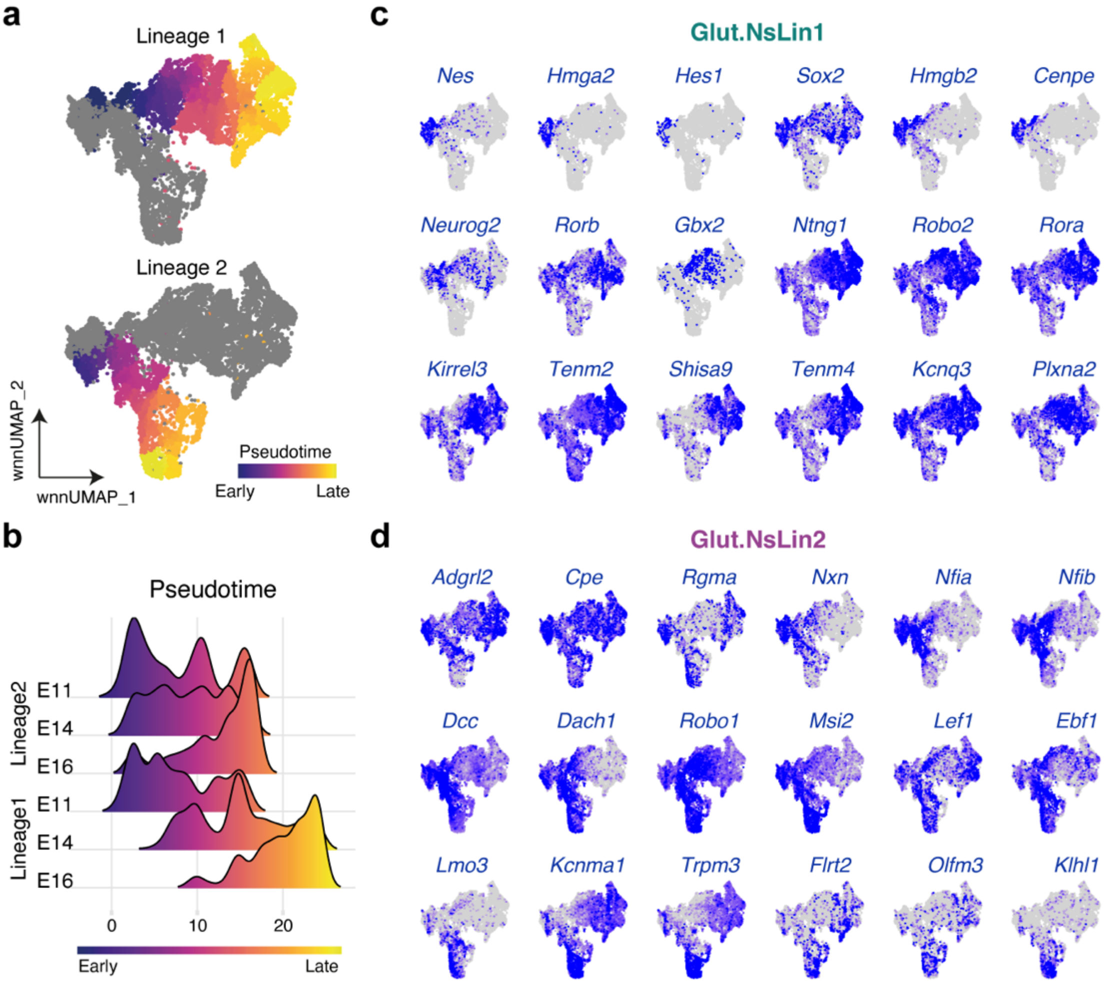


**Extended Data Fig. 5. Pseudo-temporal lineage and cell type identification of scMultiome dataset. a**, Pseudotime trajectories separated by both identified glutamatergic lineages (Top: Lineage 1; bottom: Lineage 2). **b**, Density plot displaying the population density of different lineages, colour-coded by pseudotime and separated by each time. **c**, Feature plots illustrating representative gene markers differentially expressed in neurons from lineage 1. **d**, Feature plots illustrating representative gene markers differentially expressed in neurons from lineage 2. **(Related to Fig 2)**

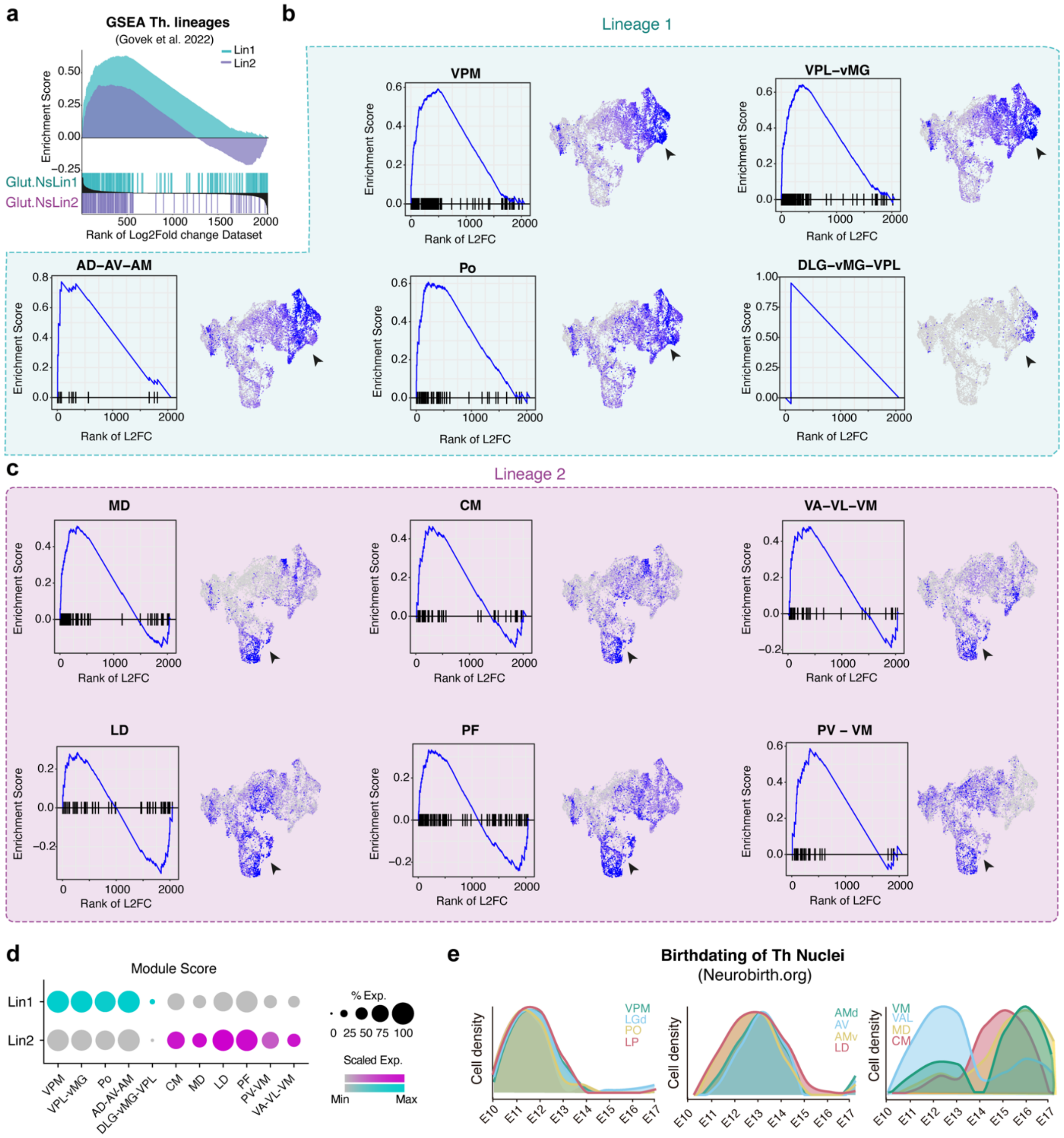


**Extended Data Fig. 6. Thalamic nuclei identification of scMultiome dataset. a**, Gene Set Enrichment Analysis (GSEA) comparing the enrichment of neurons from both lineages against the reference thalamic scRNA-seq dataset[15]. **b,** GSEA analysis (left) and feature plots (right) showing the neuronal enrichment score of thalamic nuclei identified in lineage 1 in the reference dataset. **c,** GSEA analysis (left) and feature plots (right) showing the neuronal enrichment score of thalamic nuclei identified in lineage 2 in the reference dataset. **d,** Dot plot displaying expression of thalamic nuclei module scores within each lineage. **e**, Spatio-temporal dynamic of FlashTag thalamic birthdating analysis (data from https://neurobirth.org). **(Related to Fig 2)**

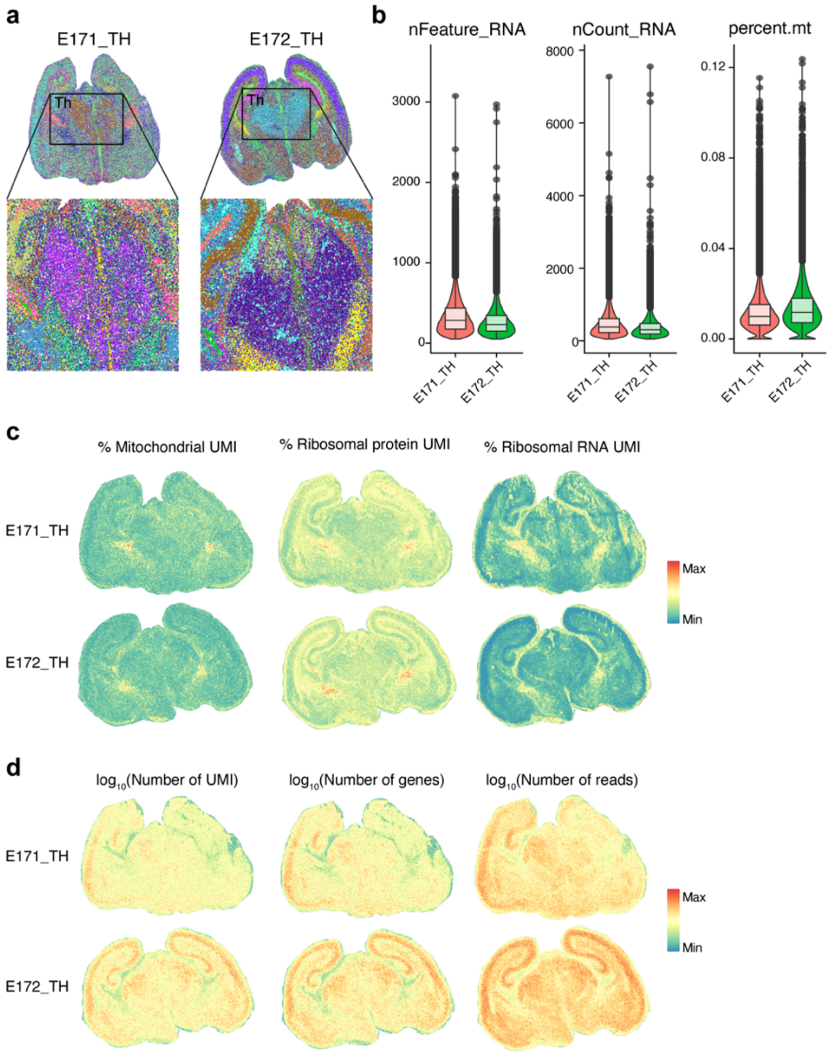


**Extended Data Fig. 7. Comprehensive analysis of whole spatial transcriptomics data. a**, Top: Slide-seq analysis of two coronal brain sections at embryonic day 17 (E17). Bottom: Inset views highlighting the thalamic region. **b**, Data quality metrics of spatial transcriptomics datasets. Violin plots display the distributions of total genes detected, total RNA reads, and the percentage of counts derived from mitochondrial genes (from left to right). **c**, Feature plots depicting the spatial distribution of the percentage of mitochondrial genes, percentage of ribosomal proteins, and percentage of ribosomal genes (from left to right). **d**, Feature plots showing the log10-transformed number of unique molecular identifiers (UMIs), log10-transformed number of genes detected, and log10-transformed number of RNA reads at spatial resolution (from left to right). **(Related to Fig. 2)**

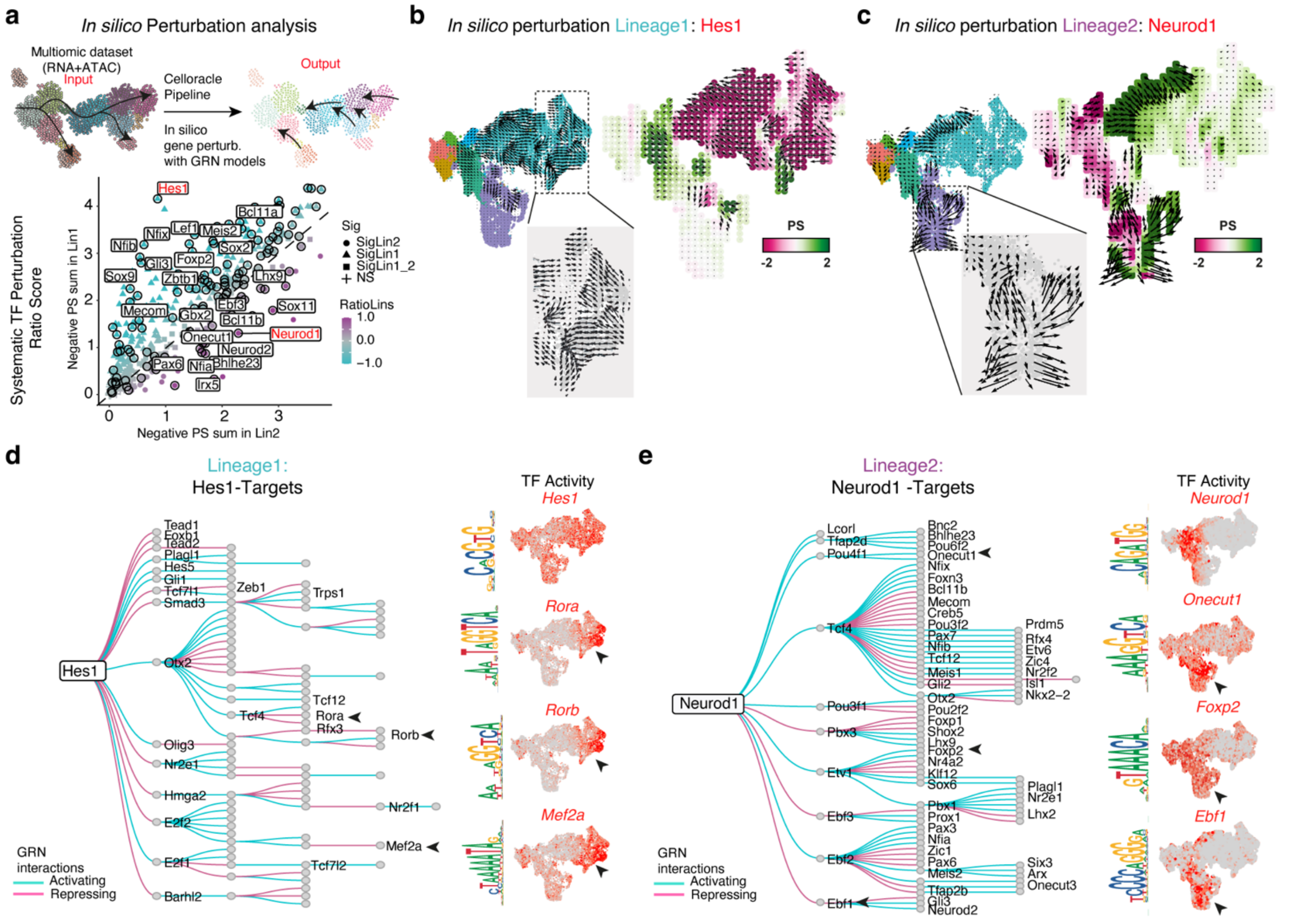


**Extended Data Fig. 8. Glutamatergic Gene Regulatory Networks (GRN) identified by single-cell multiomic enhancer-based inference. a**, Top: Schematic representation of *in silico* perturbation analysis with Celloracle. Bottom: Scatter plot shows systematic *in silico* TF simulation of negative scores for the Lineage 2 (x axis) and Lineage 1 (y axis) trajectories, respectively. **b**, Left: CellOracle vector field graphic upon UMAP embedding shows simulated vector shift after *in silico* Lineage 1: *Hes1* knockout (KO) perturbation. Right: *Hes1* KO simulation vector field with perturbation scores. **c**, CellOracle vector field graphic upon UMAP embedding shows simulated vector shift after *in silico* Lineage 2: *Neurod1* KO perturbation. Right: *Neurod1* KO simulation vector field with perturbation scores. **d**,**e**, Left: GRN subgraph for Lineage1: *Hes1* (d) and Lineage2: *Neurod1* (e) progenitors, showing first- and second-order targets. The circles represent genes for which all TFs are labelled. The edges are coloured based on TF regulatory interaction. Right: Feature plots of ChromVAR gene activity showing Specific targeted TFs for lineage 1 (d) and lineage 2 (e). **(Related to Fig 2).**

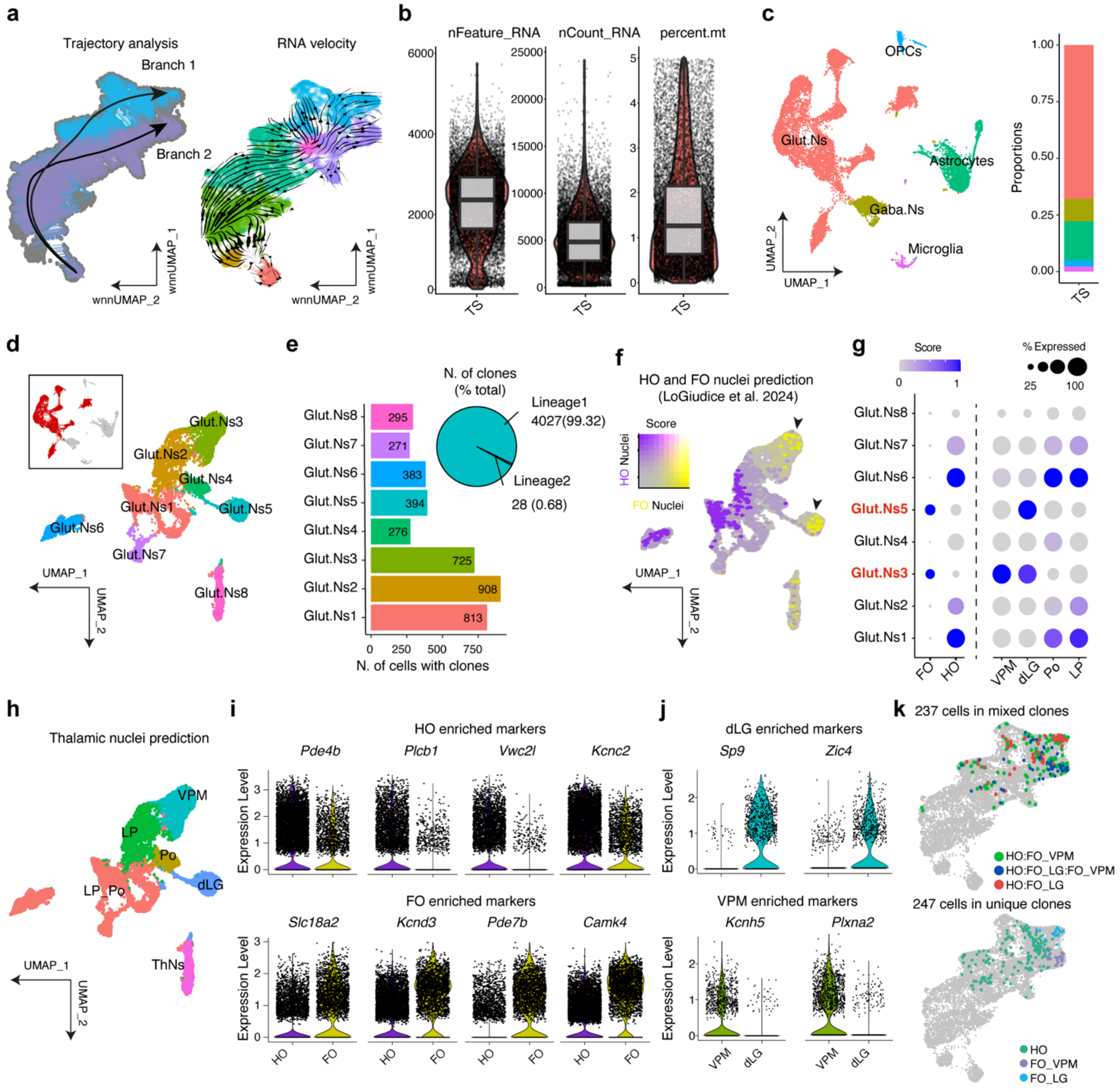


**Extended Data Fig. 9. Glutamatergic sensory lineage identified through trajectory and clonal analysis. a**, Left: UMAP visualization of inferred pseudotime trajectories from multiomic sensory lineage neurons, highlighting two isolated neuroanatomical branches. Right: RNA velocity analysis projected onto the UMAP embedding; arrow directions indicate predicted future gene expression. **b**, Data quality metrics of the TrackerSeq library at postnatal day 2 (P2). Violin plots display distributions of total genes detected, total RNA reads, and percentage of mitochondrial gene counts (from left to right). **c**, Left: UMAP visualization of single cells defined by TrackerSeq at P2. Right: Cell type proportions within the dataset. **d**, Isolation of glutamatergic neuronal clusters from (c), with glutamatergic neurons highlighted in red in the inset. **e**, Left: Number of cells assigned to cloneIDs. Right: Number of clones transferred into the multiomic dataset, split by lineage identification. **f**, UMAP representation of the TrackerSeq dataset annotated with thalamic first-order (FO) and high-order (HO) nuclei based on published data[16]. **g**, Module score assignments and average expression of FO, HO, and thalamic nuclei gene sets in the TrackerSeq dataset. h, UMAP visualization of transferred labels identifying thalamic nuclei. **i**, Violin plots showing the expression of four marker genes for HO (top) and FO (bottom) neurons. **j**, Violin plots showing the expression of two marker genes for dLG (top) and VPM (bottom) neurons. **k**, Examples of clones with sibling cells traversing mixed (top) or distinct (bottom) developmental trajectories in the multiomic UMAP embedding annotated with FO and HO labels. **(Related to Fig. 3)**

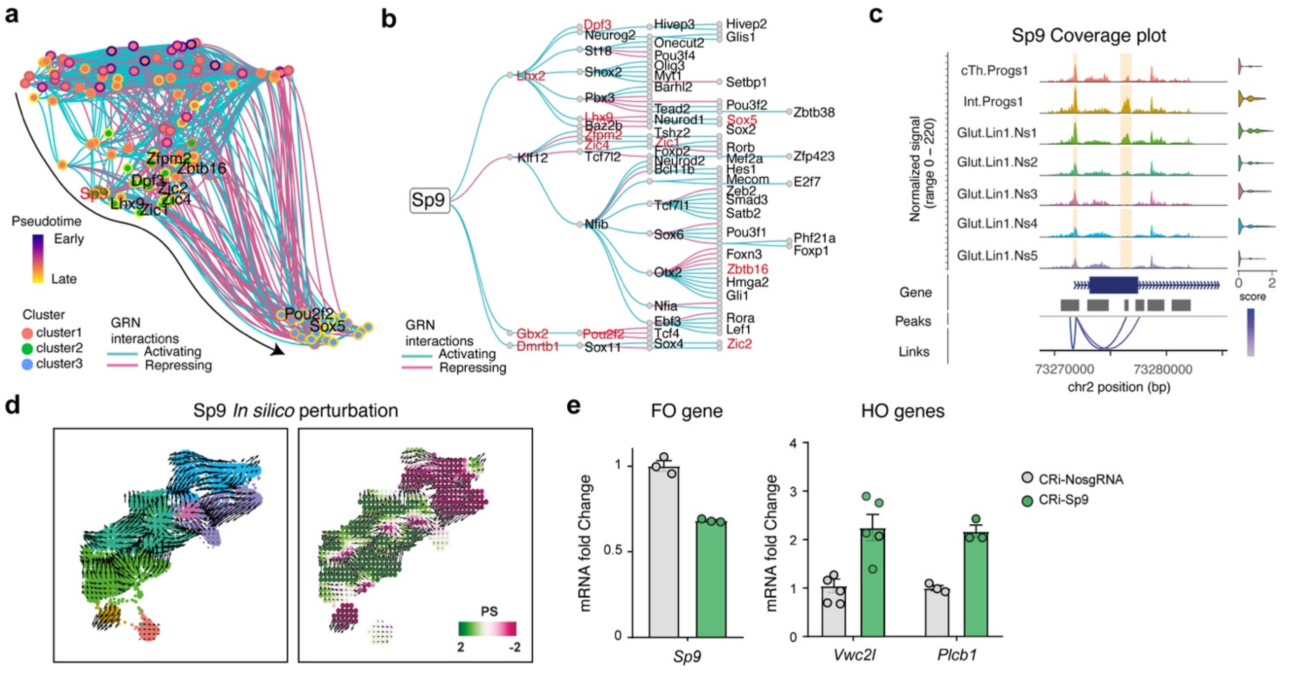


**Extended Data Fig. 10. Sp9 gene regulatory network identification and perturbation analysis.** **a**, UMAP embedding of the inferred transcription factor (TF) network in visual branch 1, based on co-expression and interaction strength. Fill colour indicates cluster identity; border colour indicates expression-weighted pseudotime; edge colour represents predicted regulatory interactions. **b**, Subgraph of the Sp9-centered gene regulatory network (GRN), highlighting first- and second-order transcriptional targets. Highlighted red TFs are specific from the visual branch trajectory. Nodes represent target genes; edge colour reflects inferred interaction strength. **c**, Genome track showing aggregated chromatin accessibility, peak-gene linkage, gene annotation, and gene expression for *Sp9* in the sensory thalamic isolation. **d**, Left: CellOracle vector field projected onto the UMAP embedding, simulating vector shifts after in silico *Sp9* knockout (KO) perturbation in branch 1. Right: *Sp9* KO simulation vector field with perturbation scores. **e**, RT–qPCR analysis showing reduced expression of *Sp9* (left) and an increased expression of high-order (HO) marker genes (right) following CRISPRi perturbation at E14.5 in primary neuronal cultures from whole thalamus ($n$ = 3-5 technical replicates). Bar graphs show the means ± SEM. **(Related to Fig. 4)**
